# Maintenance of Metabolic Plasticity Despite Relaxed Selection in a Long-Term Evolution Experiment with *Escherichia coli*

**DOI:** 10.1101/2020.06.02.130138

**Authors:** Nkrumah A. Grant, Rohan Maddamsetti, Richard E. Lenski

## Abstract

Traits that are unused in a given environment are subject to processes that tend to erode them, leading to reduced fitness in other environments. Although this general tendency is clear, we know much less about why some traits are lost while others are retained, and about the roles of mutation and selection in generating different responses. We addressed these issues by examining populations of a facultative anaerobe, *Escherichia coli*, that have evolved for >30 years in the presence of oxygen, with relaxed selection for anaerobic growth and the associated metabolic plasticity. We asked whether evolution led to the loss, improvement, or maintenance of anaerobic growth, and we analyzed gene expression and mutational datasets to understand the outcomes. We identified genomic signatures of both positive and purifying selection on aerobic-specific genes, while anaerobic-specific genes showed clear evidence of relaxed selection. We also found parallel evolution at two interacting loci that regulate anaerobic growth. We competed the ancestor and evolved clones from each population in an anoxic environment, and we found that anaerobic fitness had not decayed, despite relaxed selection. In summary, relaxed section does not necessarily reduce an organism’s fitness in other environments. Instead, the genetic architecture of the traits under relaxed selection and their correlations with traits under positive and purifying selection may sometimes determine evolutionary outcomes.

## Introduction

> “[I]f man goes on selecting, and thus augmenting, any peculiarity, he will almost certainly unconsciously modify other parts of the structure, owing to the mysterious laws of the correlation of growth.” — Charles Darwin, *On the Origin of Species*, 1859

Organisms seldom experience static conditions. Instead, they typically experience fluctuations in both their external environments and internal states. Organisms have adapted to these fluctuations by evolving a variety of mechanisms to maintain homeostasis and survive, that is to be *phenotypically robust*, in the face of environmental and genetic perturbations (Lenski et al. 2006; Frankel et al. 2010; Fraser and Schadt 2010; Siegal and Leu 2014). One mechanism to maintain homeostasis is metabolic plasticity, by which we mean the innate capacity to change metabolic fluxes in response to changes in the environment (Jia et al. 2019). Metabolic plasticity is controlled by genes, the expression of which is coupled to one or more environmental signals (Paudel and Quaranta 2019). *Bacillus subtilis,* for example, produces metabolically dormant endospores when cells are starved for nutrients (Setlow 2006). Metabolic plasticity can sometimes go awry, such as when cancer cells perform glycolysis instead of oxidative phosphorylation (Warburg 1956) to generate ATP and synthesize biomass, promoting tumorigenesis and metastatic potential (Payen et al. 2016). More generally, metabolic plasticity determines the environmental conditions in which an organism can survive and grow. However, much remains unknown about the mechanisms underlying metabolic plasticity and the resulting phenotypic robustness, and how these mechanisms and robustness have evolved and continue to evolve (Siegal and Leu 2014; Nijhout et al. 2017).

Experimental evolution with microorganisms has proven to be a powerful way to study the evolutionary process (Elena and Lenski 2003; Lenski 2017; Van den Bergh et al. 2018). These experiments typically maintain relatively simple and constant conditions, which places many traits, including metabolic plasticity, under *relaxed selection*. Accordingly, many such studies have shown losses of functions owing to antagonistic pleiotropy (including the cost of expressing unneeded traits), mutation accumulation in unused genes, or both (Cooper and Lenski 2000; Cooper et al. 2001b; Maughan et al. 2009; Leiby and Marx 2014; Lamrabet et al. 2019). However, unneeded traits may sometimes be maintained and even improved if the underlying genes serve multiple purposes, such that the unneeded trait is genetically correlated with a trait under positive selection (Bennett et al. 1990). The latter scenario has been termed *buttressing pleiotropy* (Lahti et al. 2009). Computational models and experiments with artificial organisms suggest that buttressing pleiotropy readily occurs when new functions evolve by building upon existing functions (Wagner and Mezey 2000; Lenski et al. 2003; Ostrowski et al. 2015). However, the extent of buttressing pleiotropy in biological systems remains unclear and has been little studied.

To study how metabolic plasticity evolves under relaxed selection, we analyzed both phenotypic performance and genetic changes in *Escherichia coli* populations from the long-term evolution experiment (LTEE). The 12 LTEE populations have been evolving independently in a glucose-limited minimal medium with constant aeration for more than 70,000 generations (Lenski et al. 1991; Good et al. 2017). Samples are frozen every 500 generations, generating a “frozen fossil record” from which bacteria can be revived for genetic and phenotypic analyses, including measuring their fitness in the LTEE environment and other environments that differ in various respects. By 50,000 generations the populations were, on average, about 70% more fit than their common ancestor in the LTEE environment (Wiser et al. 2013). Most of that improvement occurred in the first 10,000 generations, but the populations have continued to improve at slower rates throughout the experiment (Lenski et al. 2015; Lenski 2017). The bacteria experienced relaxed selection for anaerobic growth during the long duration of their evolution in the strictly oxic LTEE environment. To date, several hundred clones and more than 1400 mixed population samples have been sequenced (Tenaillon et al. 2016; Good et al. 2017), providing material for examining the coupling between aerobic and anaerobic fitness and the underlying genetic changes. Six LTEE populations evolved hypermutability during the experiment (Tenaillon et al. 2016; Good et al. 2017), which should increase the rate at which unused genes accumulate mutations and unused functions decay over time (Cooper and Lenski 2000; Leiby and Marx 2014). The hypermutable lineages exhibited ~100-fold increases in their point-mutation rate (Sniegowski et al. 1997; Wielgoss et al. 2013), but the rate of fitness gain in these lineages increased by only a few percent relative to other populations (Wiser et al. 2013; Lenski et al. 2015). This difference facilitates disentangling the effects of antagonistic pleiotropy from those of mutation accumulation on the phenotypic and genomic consequences of relaxed selection.

In addition to the strength of the LTEE model system for asking evolutionary questions, *E. coli* is an excellent model for studying the evolution of metabolic plasticity. As a facultative anaerobe, *E. coli* is able to survive, grow, and reproduce in both oxic and anoxic environments (Unden et al. 1994). This plasticity allows *E. coli* to inhabit diverse environments that vary in oxygen availability, from the gastrointestinal tracts of mammals (and some birds and reptiles) to freshwater and soil environments (Gordon and Cowling 2003). The molecular control of this plasticity is well-understood, and two global regulatory systems play critical roles (Gunsalus and Park 1994; Unden and Bongaerts 1997). The Fumarate and Nitrate Reductase protein (FNR), encoded by the *fnr* gene, is a transcription factor that directly senses oxygen (Gunsalus and Park 1994; Kang et al. 2005). The Anoxic Respiratory Control system, encoded by *arcA* and *arcB*, is a canonical two-component regulatory system that responds to oxygen and the redox status of the cell, as shown in Figure 1 (Iuchi and Lin 1988; Gunsalus and Park 1994; Unden and Bongaerts 1997). The transcriptional control conferred by the *fnr* and *arcAB* regulons allows *E. coli* cells to commit physiologically to either aerobic or anaerobic metabolism, depending on oxygen availability. Mutations in either regulon may disrupt this control (Melville and Gunsalus 1990). In particular, some mutations in *arcAB* have been shown to alter metabolism by causing the constitutive expression of genes that would normally be responsive to oxygen concentration and internal redox balance (Iuchi and Lin 1988; Saxer et al. 2014).

**Figure 1:**
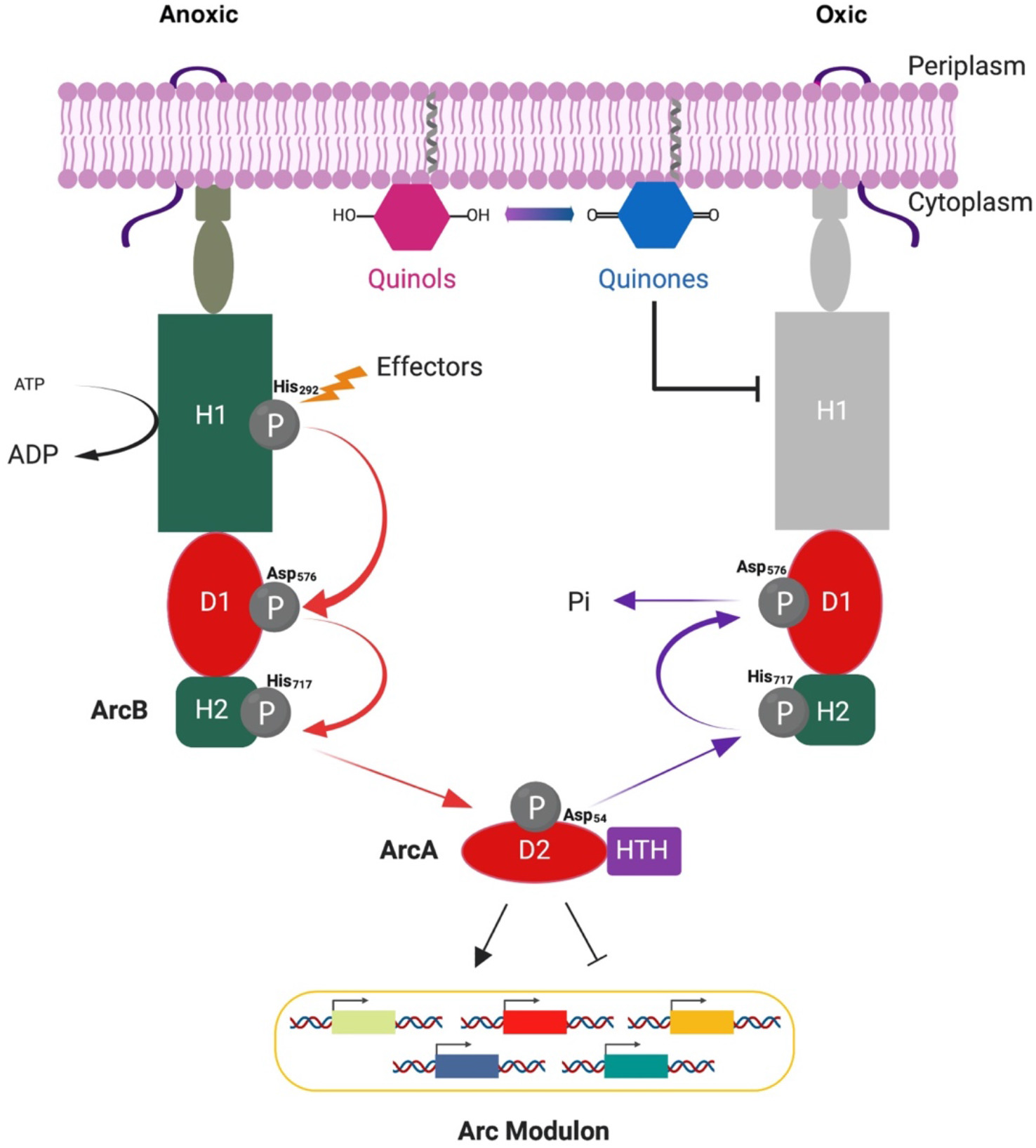
Schematic representation of ArcAB-dependent gene regulation. In an anoxic environment, the transmembrane sensor kinase ArcB undergoes autophosphorylation. This reaction is enhanced by fermentation metabolites such as D-lactate, pyruvate and acetate that act as effectors. Three conserved residues (His_292_, Asp_576_, His_717_) in ArcB sequentially transfer the phosphoryl group onto ArcA, the response regulator, at a conserved Asp54 residue. Phosphorylated ArcA (ArcA-P), in turn, represses the transcription of many operons involved in respiratory metabolism, while activating those encoding proteins involved in fermentative metabolism. In an oxic environment, ArcB autophosphorylation ceases and ArcA is dephosphorylated by reversing the phosphorelay, leading to the release of inorganic phosphate into the cytoplasm. Figure adapted from Kwon et al. (2003).

In this study, we sought to determine whether relaxed selection during 50,000 generations of strictly aerobic growth led to the loss of anaerobic performance and the associated metabolic plasticity. Alternatively, genetic correlations between aerobic and anaerobic physiology might have favored the maintenance or even improvement of anaerobic metabolism during evolution in the oxic environment of the LTEE. This alternative, if observed, might reflect the fact that anaerobic metabolism evolved more than 2 billion years before the origin of aerobic metabolism (Müller 1977; Soo et al. 2017), such that the genetic and biochemical networks underpinning metabolism in these conditions might be tightly coupled. It could also indicate the biochemical promiscuity of many proteins involved in metabolism (Nam et al. 2012). In any case, we tested the phenotypic correlation in performance by measuring the fitness of the evolved LTEE clones against a marked ancestor in oxic and anoxic environments. In fact, anaerobic growth capacity was not only maintained under relaxed selection, but in some cases actually improved. Fitness gains were seen even in some populations that evolved high mutation rates—a change that promoted mutation accumulation in anaerobic-specific genes—and despite mutations affecting the ArcAB regulon.

Our results highlight the importance of understanding how a trait is encoded in a genetic network, and how that encoding affects its evolutionary fate in the absence of direct selection. A better understanding of the mechanisms that maintain anaerobic growth may also help synthetic biologists design more robust systems by exploiting pleiotropy (Stirling et al. 2017; Blazejewski et al. 2019; Geng et al. 2019). In addition, this work may provide a framework for better predicting how the genetic encoding of traits affects an organism’s evolutionary potential, including in response to ecological challenges such as those caused by climate change.

## Materials and Methods

### Long-Term Evolution Experiment

The LTEE consists of 12 *E. coli* populations derived from a common ancestral strain, REL606 (Lenski et al. 1991). Six populations descend directly from REL606. The other six descend from REL607, which differs from REL606 by two mutations that are selectively neutral under LTEE conditions (Tenaillon et al. 2016). One is a point mutation in the *araA* gene that allows REL607 to utilize arabinose, and the second is an inadvertent secondary mutation of no consequence. The mutation in *araA* provides a phenotypic marker that can be readily scored in the competition assays used to measure relative fitness. When plated on tetrazolium arabinose (TA) indicator agar plates, REL606 and its direct descendants form red colonies, whereas REL607 and its descendants form white colonies. The ability to freeze and revive viable strains has allowed the establishment of the frozen fossil record that includes samples from all 12 populations at 500-generation intervals. This record allows both genotypic and phenotypic changes to be quantified retrospectively. In this study, we examined ancestral and evolved clones that were frozen at generations 2,000, 10,000 and 50,000 (supplemental PDF, Table S1).

### Culture Conditions

Unless noted otherwise, we grew strains in oxic and anoxic environments in 10 mL of Davis Mingioli minimal salts medium supplemented with 25 μg/mL glucose (DM25). We prepared anaerobic media by boiling 500-mL batches for 25 min while sparging in nitrogen gas using a hypodermic needle inserted through a butyl rubber stopper. Cultures were incubated at 37°C in 50-mL Erlenmeyer flasks, with orbital shaking at 120 rpm in the oxic but not the anoxic environment. These conditions are the same as those used during the LTEE, except for the absence of oxygen and shaking during anaerobic growth.

### Competition Assays

We revived frozen clones isolated from each LTEE population at generations 2,000, 10,000 and 50,000. We used clones for which whole-genome sequences are available (Tenaillon et al. 2016). The notation Ara-1 to Ara-6 denotes clones descended from the ancestral strain REL606, while Ara+1 to Ara+6 are derived from REL607. We excluded from these assays the 50,000-generation clones from three populations (Ara-2, Ara-3, Ara+6), as their evolved phenotypes make the assays unreliable (Wiser et al. 2013). We also excluded the clones sampled from Ara+6 at both earlier time-points because their growth was erratic in the anoxic environment. Clones were revived by inoculating 15 μL of thawed frozen stock into 10 mL of Luria-Bertani (LB) broth, and they were grown at 37°C in an orbital shaker under atmospheric conditions for 24 h. We then diluted each competitor 1:10,000 into DM25 medium for a preconditioning step that depended on whether the assay would be performed in the oxic or anoxic environment. For the former, the competitors were preconditioned in DM25 under the standard LTEE conditions. For the latter, the competitors were preconditioned in anaerobic DM25 in an anaerobic chamber under a 95%-N_2_:5%-H_2_ atmosphere. Each preconditioned competitor was then diluted 1:200 into a flask containing the relevant medium under the appropriate atmosphere, and the competition ran for one day, during which time the combined population grew 100-fold. The one-day competition assays encompassed the same lag, exponential growth, and stationary phases as populations experienced during the LTEE (Lenski et al. 1991; Vasi et al. 1994). We competed the Ara^−^ evolved clones against REL607 and the Ara^+^ evolved clones against REL606. Competitions were replicated fivefold for all clones across all generations and in both environments. Owing to technical errors, only four replicates yielded data for: Ara-1 at 2,000 generations, and Ara+1 and Ara+4 at 50,000 generations, in the oxic environment; and Ara-2 and Ara+2 at 2,000 generations, Ara+1 at 10,000 generations, and Ara-1 at 50,000 generations in the anoxic environment. We calculated the fitness of an evolved clone relative to the ancestral competitor as the ratio of their growth rates realized during the competition assay (Lenski et al. 1991; Wiser et al. 2013).

### Genomic and Statistical Analyses

The genomes of the clones used in this study were previously sequenced (Tenaillon et al. 2016). We used an online tool (http://barricklab.org/shiny/LTEE-Ecoli/) to identify all of the mutations discovered specifically in the *fnr* and *arcAB* genes. To identify genes regulated in response to oxygen levels, Salmon et al. (2005) performed a Bayesian analysis of gene-expression data obtained for *E. coli* K12 in oxic and anoxic environments. We then used the set of genes from that study that showed differential expression between oxic and anoxic conditions at a posterior probability greater than 99%. Those genes were mapped onto the REL606 reference genome using the OMA (Altenhoff et al. 2018) and EcoCyc (Keseler et al. 2017) databases. We call genes that are upregulated under oxic conditions “aerobic-specific genes,” and those upregulated under anoxic conditions “anaerobic-specific genes.” We performed binomial tests to compare the numbers of mutations in the LTEE-derived genomes in aerobic- and anaerobic-specific genes to a null expectation based on the summed length of genes in the two gene sets. That analysis was conducted using an R script called aerobic-anaerobic-genomics.R.

In addition, the LTEE metagenomics dataset includes mutations found by sequencing whole-population samples for all 12 populations through 60,000 generations. These mutations were downloaded from https://github.com/benjaminhgood/LTEE-metagenomic/. The data were reformatted (*.csv) and analyzed using an R script called aerobic-anaerobic-metagenomics.R. In brief, the cumulative number of mutations observed in each population was plotted, after normalizing by gene length, for various categories of mutations. In particular, we examined subsets of these data based on mutation type (nonsynonymous, synonymous, and all others including indels, nonsense, and structural variants) and by function (occurring in the aerobic- or anaerobic-specific genes). To generate a null expectation, we chose 10,000 random sets of genes (with the same cardinality as the aerobic- and anaerobic-specific genes in the specific comparison), and the cumulative number of mutations in each set was calculated. The proportion of replicates in which the cumulative number of mutations in the random set was larger than the corresponding number in the aerobic-or anaerobic-specific gene set was used as an empirical *p*-value for testing statistical significance. For visualization, the relevant figures show only the middle 95% of the cumulative mutations for 1,000 (rather than 10,000) random sets. Statistical analyses were performed in R (version 3.5.0; 2018-04-23). Datasets and R analysis scripts are available on the Dryad Digital Repository (DOI pending publication).

### Aerobic and Anaerobic Network Connectivity

We used the Cytoscape platform (Shannon et al. 2003) to visualize the biomolecular interaction network between aerobic- and anaerobic-specific genes. In brief, we imported our gene sets into Cytoscape and then used the software to query the STRING database (Szklarczyk et al. 2017), which includes empirically known and computationally predicted protein-protein interactions. Interactions are evaluated using seven lines of evidence, and a score is assigned to each (Szklarczyk et al. 2017). The STRING software then computes a combined “confidence score” for each interaction, which is effectively the likelihood that the interaction truly exists given the evidence. We constructed our network of the aerobic- and anaerobic-specific genes showing only those interactions with confidence scores greater than 70%, a threshold considered “high” by the database curators.

## Results

### Signatures of Selection on Aerobic- and Anaerobic-specific Genes in the LTEE

We hypothesized that a subset of the aerobic-specific genes experienced positive selection to acquire mutations that better adapt the bacteria to the LTEE environment. By contrast, we expect that many anaerobic-specific genes were under relaxed selection in the LTEE. We tested these predictions using the genomic and metagenomic datasets spanning 50,000 and 60,000 generations, respectively (Tenaillon et al. 2016; Good et al. 2017). At various times, 6 of the 12 LTEE populations evolved roughly 100-fold higher point-mutation rates than the ancestral strain (Sniegowski et al. 1997; Tenaillon et al. 2016; Good et al. 2017). These mutator populations gained fitness slightly faster than the non-mutator populations (Wiser et al. 2013; Lenski et al. 2015). However, genomic evolution in these populations was dominated by the accumulation of random mutations (Tenaillon et al. 2016; Couce et al. 2017; Maddamsetti et al. 2017). For these reasons, we made and tested separate predictions for the mutator and non-mutator populations. For the nonmutator populations, where previous studies found compelling evidence for positive selection (Woods et al. 2006; Tenaillon et al. 2016; Good et al. 2017), we predicted that aerobic-specific genes would have more mutations than anaerobic-specific genes. By contrast, in the mutator populations, previous studies indicated that random mutations (neutral or nearly neutral) accumulated in those genes under relaxed selection, given the high mutation pressure. Of course, some sites even within aerobic-specific genes could have accumulated neutral or nearly neutral mutations in the oxic LTEE environment. However, purifying selection should lead to fewer mutations in aerobic-than in anaerobic-specific genes in the mutator populations.

To test these predictions, we examined the sets of 345 and 227 anaerobic and aerobic-specific genes, respectively, as described in the Materials and Methods (supplemental file 1). Given 4,143 protein-coding genes in the genome of the LTEE ancestor (Jeong et al. 2009), the anaerobic- and aerobic-specific genes constitute 8.3% and 5.5% of that total, respectively. We asked whether these two sets accumulated different numbers of mutations in clones sampled from the LTEE populations at 50,000 generations. We controlled for differences in mutational target size by summing over the length of the genes in each set. In clones from the six non-mutator lineages, the aerobic-specific genes had 38 mutations, whereas the anaerobic-specific genes had 21 mutations (two-tailed binomial test: *p* < 10^−6^). By contrast, the hypermutator clones had 836 mutations in anaerobic-specific genes and 333 mutations in aerobic-specific genes (two-tailed binomial test: *p* = 0.0040). The first result is consistent with stronger positive selection for beneficial mutations in aerobic-than anaerobic-specific genes. The second result is consistent with relaxed selection on anaerobic-specific genes, which could also be described as stronger purifying selection on aerobic-specific genes.

Next, we examined these dynamics using whole-population metagenomic data that includes mutations that fixed as well as those that reached a frequency above ~5% in a population through ~60,000 generations (Good et al. 2017). Rather than reanalyzing these data from scratch, we analyzed the dataset previously generated by Good et al. (2017). We first visualized the evolutionary dynamics for all mutations in aerobic- and anaerobic-specific genes in each population (supplemental PDF, fig. S1). For the non-mutator populations (panels A–F in fig. S1), mutations in aerobic-specific genes were more common than those in anaerobic-specific genes, especially during the first 10,000 generations, indicating that many mutations in aerobic-specific genes experienced positive selection. In the populations that evolved hypermutator phenotypes (panels G–L in fig. S1), however, it is difficult to tell by eye whether the rate of molecular evolution differs between these two sets of genes. Therefore, we counted the number of observed mutations in aerobic and anaerobic-specific genes in each population over time, normalized by gene length.

When we examine the occurrence of nonsynonymous mutations, all six populations that were never mutators, along with two others (Ara-1, Ara-3) before they became hypermutable, have substantially more mutations in aerobic-specific genes than expected under the null distribution calculated by resampling random sets of 227 genes (the cardinality of the aerobic-specific genes) (fig. 2). The probability of this directional outcome occurring by chance under a one-tailed binomial expectation is (1/2)^8^ = 1/256 ≈ 0.004, which is thus very significant. The number of nonsynonymous mutations in anaerobic-specific genes in those same populations, by contrast, is much lower in every case. For the mutator populations, including the two (Ara-1, Ara-3) that evolved hypermutability fairly late in the LTEE, the rates of mutations in anaerobic- and aerobic-specific genes track one another more closely. In at least three of these populations (Ara-2, Ara+3, Ara+6), the rates of mutation accumulation decreased later in the LTEE. These decelerations correspond to reversions or compensatory alleles that arose in those populations and caused their mutation rates to decline (Tenaillon et al. 2016; Good et al. 2017). Some of these populations also suggest a slower rate of mutation accumulation in aerobic-than in anaerobic-specific genes, in particular near the end of the time course. Indeed, two mutator populations, Ara+3 and Ara+6, accumulated significantly fewer nonsynonymous mutations in aerobic genes than expected under the null distribution (non-parametric bootstrap with 10,000 replicates, *p* < 0.0001 for each). The slower mutation accumulation in aerobic-specific genes suggests purifying selection, as expected because the mutator phenotype increases the rate of deleterious mutations.

**Figure 2:**
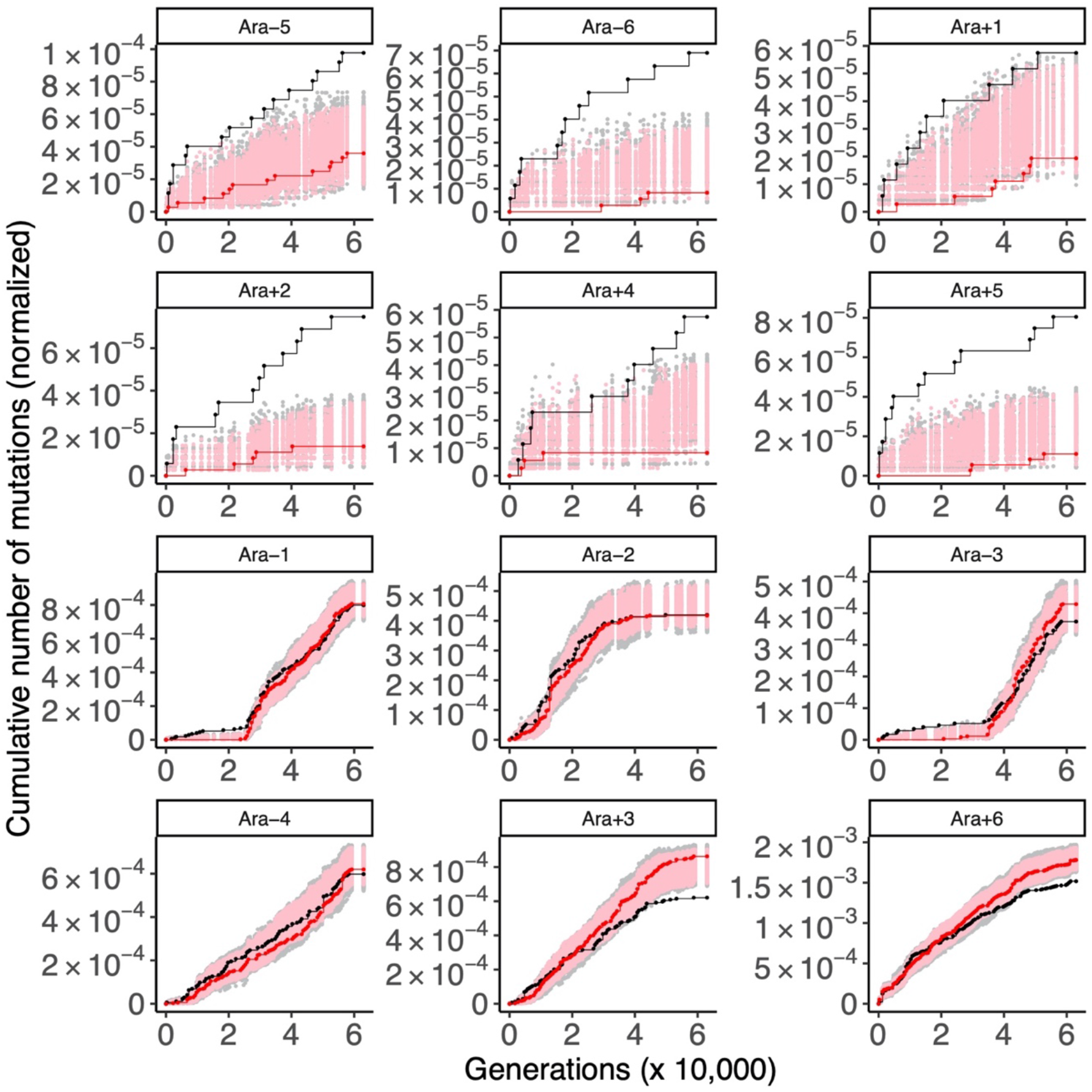
Cumulative numbers of mutations in aerobic- and anaerobic-specific genes in the LTEE whole-population samples. Each panel shows the number of nonsynonymous mutations in aerobic-(black) and anaerobic-specific (red) genes in the indicated population through 60,000 generations. For comparison, random sets of genes of equal cardinality to the aerobic-(227) or anaerobic-specific (345) gene sets were sampled 1,000 times, and the cumulative number of nonsynonymous mutations was calculated to generate a null distribution of the expected number of mutations for each population. The gray and pink points show 95% of these null distributions (excluding 2.5% in each tail) for aerobic and anaerobic comparisons, respectively.

It might also reflect, in part, saturation of possible beneficial mutations in aerobic-specific genes, in accord with a “coupon-collecting” model of molecular evolution (Good et al. 2017). Population Ara-4 is an outlier, however, in that its rate of mutation accumulation in aerobic-specific genes slightly exceeded the rate observed in anaerobic-specific genes for most of its history, despite its mutator phenotype (fig. 2).

To look more deeply into the role of purifying selection, we examined the accumulation of insertions and deletions (indels), structural variants (including those generated by transposable elements), and nonsense mutations in protein-coding genes in all 12 populations. These types of mutations typically destroy protein function. Although such knockout mutations are sometimes beneficial in evolution experiments (e.g., Cooper et al. 2001b), they would be highly deleterious in conserved genes under purifying selection as well as in genes under positive selection to finetune protein function (Maddamsetti et al. 2017). If aerobic-specific genes faced strong purifying selection in the mutator populations, we reasoned that indels, structural variants, and nonsense mutations would be underrepresented in them (fig. 3). Indeed, that was the case in four of the six hypermutator populations (nonparametric bootstrap with 10,000 replicates: *p* = 0.0003 for Ara-3; *p* = 0.0014 for Ara-4; *p* < 0.0001 for Ara+3; *p* = 0.0003 for Ara+6). On balance, these observations indicate stronger purifying selection on aerobic-than on anaerobic-specific genes in the LTEE, especially in the populations that evolved hypermutable phenotypes. This finding is consistent with the later evolution of anti-mutator alleles that reduced or reverted mutation rates to the ancestral level in most of the populations that evolved hypermutability. The fact that mutations accumulated more slowly in anaerobic-specific genes than expected under the null distribution in two mutator populations (nonparametric bootstrap with 10,000 replicates: *p* < 0.0001 for Ara+3; *p* = 0.0035 for Ara+6) suggests that some anaerobic-specific genes might also have experienced purifying selection, indicating functionality even during aerobic growth.

**Figure 3:**
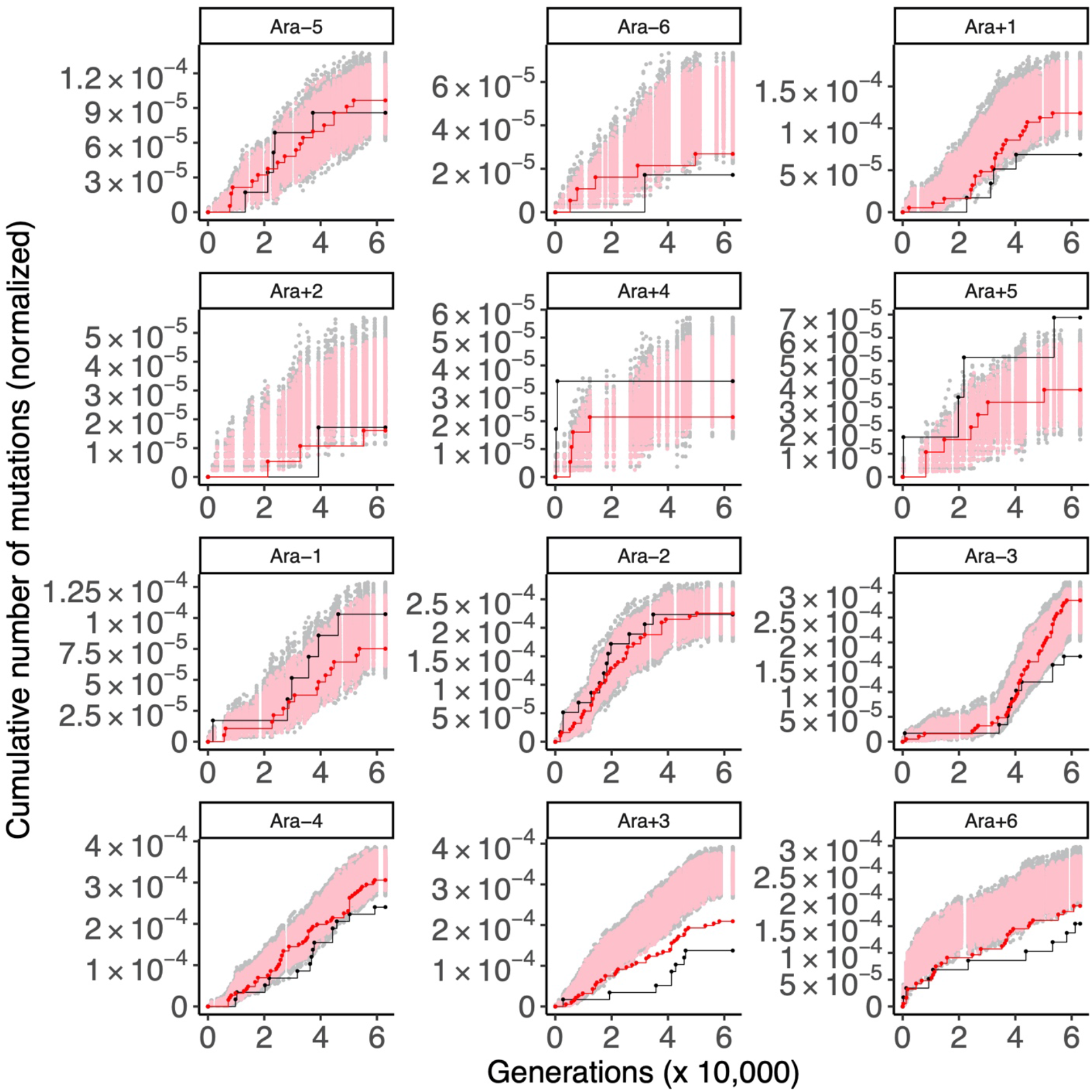
Cumulative numbers of indel, nonsense, and structural mutations in protein-coding genes in the LTEE whole-population samples. The black and red points are mutations in aerobic- and anaerobic-specific genes, respectively. The gray and pink points show the corresponding null distributions based on randomized gene sets. See Figure 2 for additional details.

For synonymous mutations, which are effectively neutral in the vast majority of cases, we expect to see many more of them in the mutator populations, and indeed that is the case. We also do not expect to see any systematic association with aerobic-or anaerobic-specific genes. That expectation is also fulfilled: three of the six mutator populations had more synonymous mutations in aerobic-than in anaerobic-specific genes, and the other three show the opposite trend (supplemental PDF, fig. S2). It is a bit puzzling that the difference between the two gene sets is so noticeable in some cases in one direction or the other. These differences might reflect hitchhiking, whereby several synonymous mutations affecting one or the other gene set, all on the same background, were pushed to high frequency (or pulled to extinction) in a particular population (Maddamsetti et al. 2015). In any case, there is no overall pattern across the six mutator populations for synonymous mutations (supplemental PDF, fig. S2).

### Mutations in Genes that Regulate Metabolic Plasticity in Response to Oxygen

After establishing the genome-wide signatures of selection on aerobic- and anaerobic-specific genes, we now turn our attention to three particular genes known to regulate metabolic plasticity in response to oxygen availability: *fnr*, *arcA*, and *arcB* (fig. 1). Only two LTEE populations, both mutators (Ara+3, Ara+6), have nonsynonymous mutations in the *fnr* gene; in both populations, the mutations arose well after the populations had evolved hypermutability. By contrast, 11 of the 12 populations have nonsynonymous mutations in *arcA, arcB*, or both (fig. 4A). The other population (Ara-6) has a 9-bp deletion in *arcA.* In another population, Ara-2, two lineages designated S and L have coexisted since about generation 6,000 (Rozen et al. 2005), and only the S lineage has a mutation in *arcA*. Many of the mutations in *arcA* and *arcB* were already present in the clones sequenced at 10,000 generations (fig. 4A), and *arcA* was previously identified as showing a signature of strong positive selection in the LTEE (Tenaillon et al. 2016). We mapped the *arcA* and *arcB* mutations in the 50,000-generation clones onto the encoded protein structures. Mutations in *arcA* impact both the response regulator and DNA binding domains of the protein (fig. 4B), and mutations in *arcB* map to several protein domains including the histidine kinase and histidine kinase receptor (fig. 4C). There are several ways that these mutations might affect metabolic plasticity in the evolved bacteria. Mutations in either gene could affect the stability of the proteins or their activities, thereby (i) altering the capacity of ArcB to sense redox changes through the quinone pool; (ii) altering the rate or efficiency of phosphoryl transfer between the ArcB protein domains; (iii) decreasing the extent of ArcA phosphorylation; (iv) increasing the rate of ArcA dephosphorylation; (v) decreasing the extent of ArcA phosphorylation-dependent oligomerization; or (vi) altering the DNA binding efficiency of ArcA. In any case, these mutations may impact ArcAB signaling and might thus affect the ability of the evolved strains to grow in an anoxic environment.

**Figure 4:**
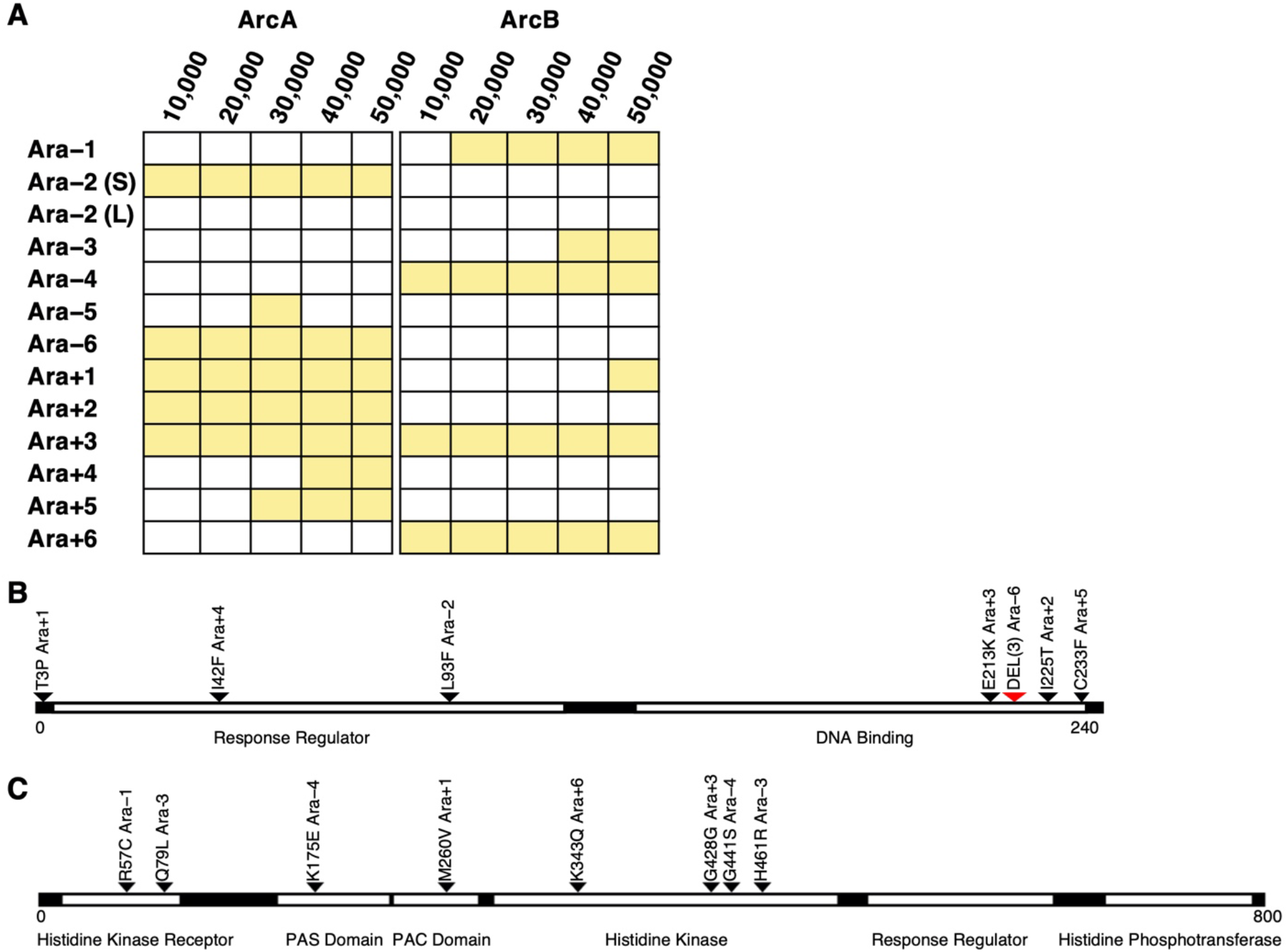
Mutations in *arcA* and *arcB* in clones from the LTEE. (A) Shading indicates that the clone has a mutation in *arcA* or *arcB*, which encode proteins ArcA and ArcB, respectively. Clones from two coexisting lineages, labeled S and L, are shown for population Ara–2. Sequence data are from Tenaillon et al. (2016), with one exception: Plucain et al. (2014) show that the *arcA* mutation had already fixed in the Ara–2 S lineage by 10,000 generations. Mutations present in the 50,000-generation clones are mapped onto the functional domains of the (B) ArcA and (C) ArcB proteins. The red arrow marks a deletion of 3 amino acids; all other mutations are point mutations resulting in amino-acid substitutions.

### Fitness of Evolved Bacteria Under Oxic and Anoxic Conditions

Half of the LTEE populations were founded by *E. coli* B strain REL606 and half by REL607, an *araA* mutant of REL606. This mutation is selectively neutral in the LTEE environment (Lenski et al. 1991; Wiser et al. 2013), and it provides a readily scored marker for distinguishing competitors in assays of relative fitness. However, it was unknown whether the *araA* mutation is also neutral under anoxic conditions. To that end, we competed REL606 and REL607 in oxic and anoxic environments, with other conditions the same as those used in the LTEE. We saw no significant differences in relative fitness in either the oxic (*t* = 0.6972, d.f. = 4, two-tailed *p* = 0.5241) or anoxic (*t* =1.1433, d.f. = 4, two-tailed *p* = 0.3167) environment. Thus, the *araA* mutation can serve as a useful marker for assaying the relative fitness of evolved and ancestral clones in both the anoxic and oxic environments.

We expected that the evolved bacteria would be better adapted to the oxic environment, where they evolved, than to the anoxic environment. To test this hypothesis, we competed clones sampled at 2,000, 10,000, and 50,000 generations against the reciprocally marked ancestral strains. As explained in the Materials and Methods section, we excluded one population (Ara+6) at all three time points, and two others (Ara-2, Ara-3) at the last time point, because of technical difficulties associated with enumerating these competitors. We calculated fitness as the ratio of the realized growth rate of the evolved clone relative to that of the ancestor during a competition. This metric integrates the effects of differences in lag, growth, and stationary phases (Lenski et al. 1991; Vasi et al. 1994). Figure 5 shows the results obtained for each evolved clone, with standard errors based on replicate competition assays for that clone. Note that all points lie below the isocline corresponding to equal fitness in the anoxic and oxic environments, consistent with our hypothesis. As an overall assessment, we performed paired comparisons of fitness in the two environments at each time point, and in all cases the difference was highly significant (paired *t*-tests; generation 2,000: *t* = 8.7849, df = 10, one-tailed *p* < 0.0001; generation 10,000: *t* = 8.7986, df = 10, one-tailed *p* < 0.0001; generation 50,000: *t* = 5.9556, df = 8, one-tailed *p* = 0.0002). Indeed, all 31 clones tested had higher estimated fitness values in the oxic environment than in the anoxic one (sign test, *p* << 0.0001). Figure 6 shows the grand mean fitness values in the two environments over time, with confidence limits based on the replicate populations.

**Figure 5:**
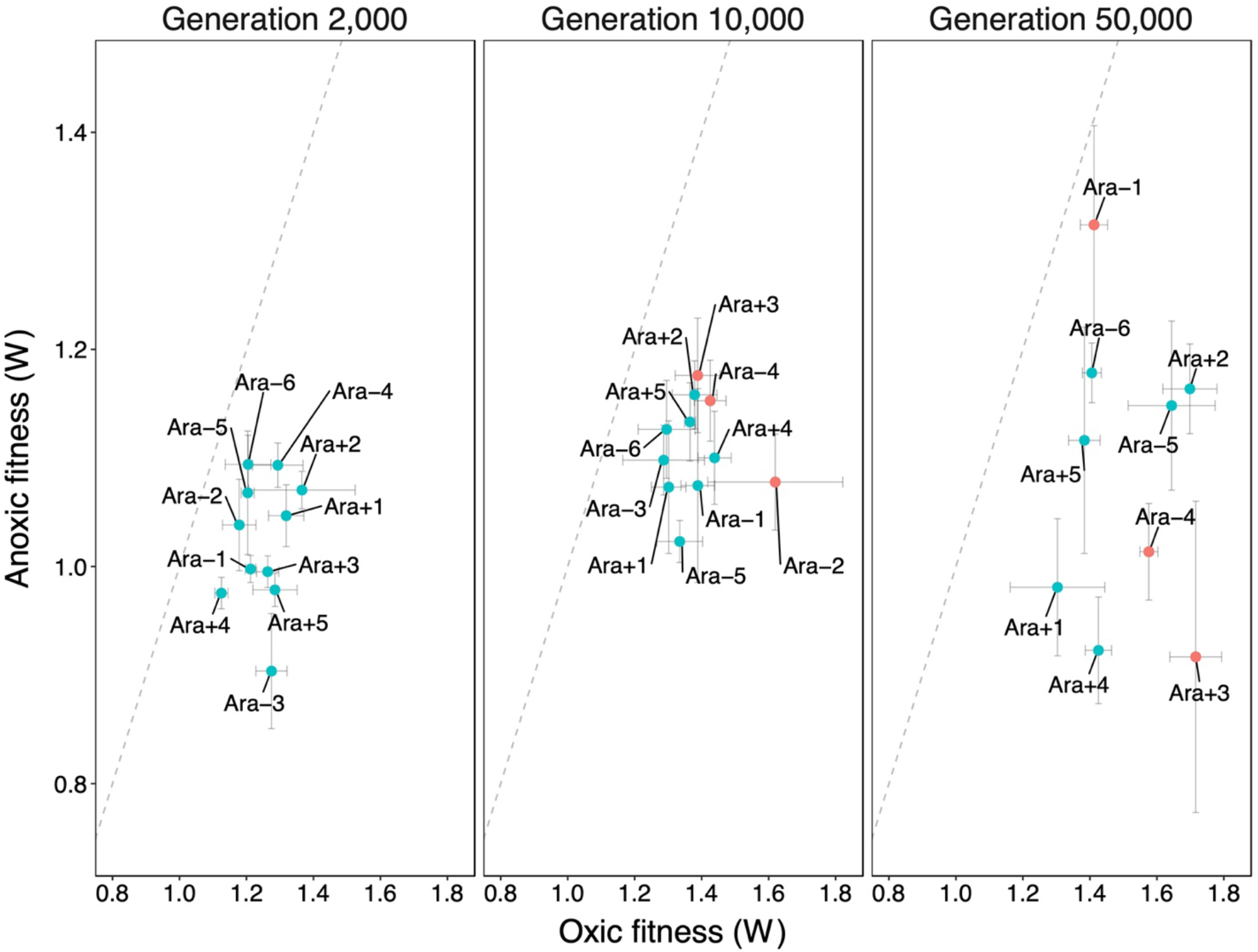
Evolved clones are better adapted to the aerobic environment in which they evolved than to the novel, but otherwise identical, anoxic environment. Each point is the mean fitness for an evolved clone sampled from the indicated population at three different generations, measured relative to the LTEE ancestor with the opposite marker state. Orange and teal points indicate that the corresponding population had or had not evolved hypermutability, respectively. Error bars show the standard errors based on replicated fitness assays in each environment for each clone.

**Figure 6:**
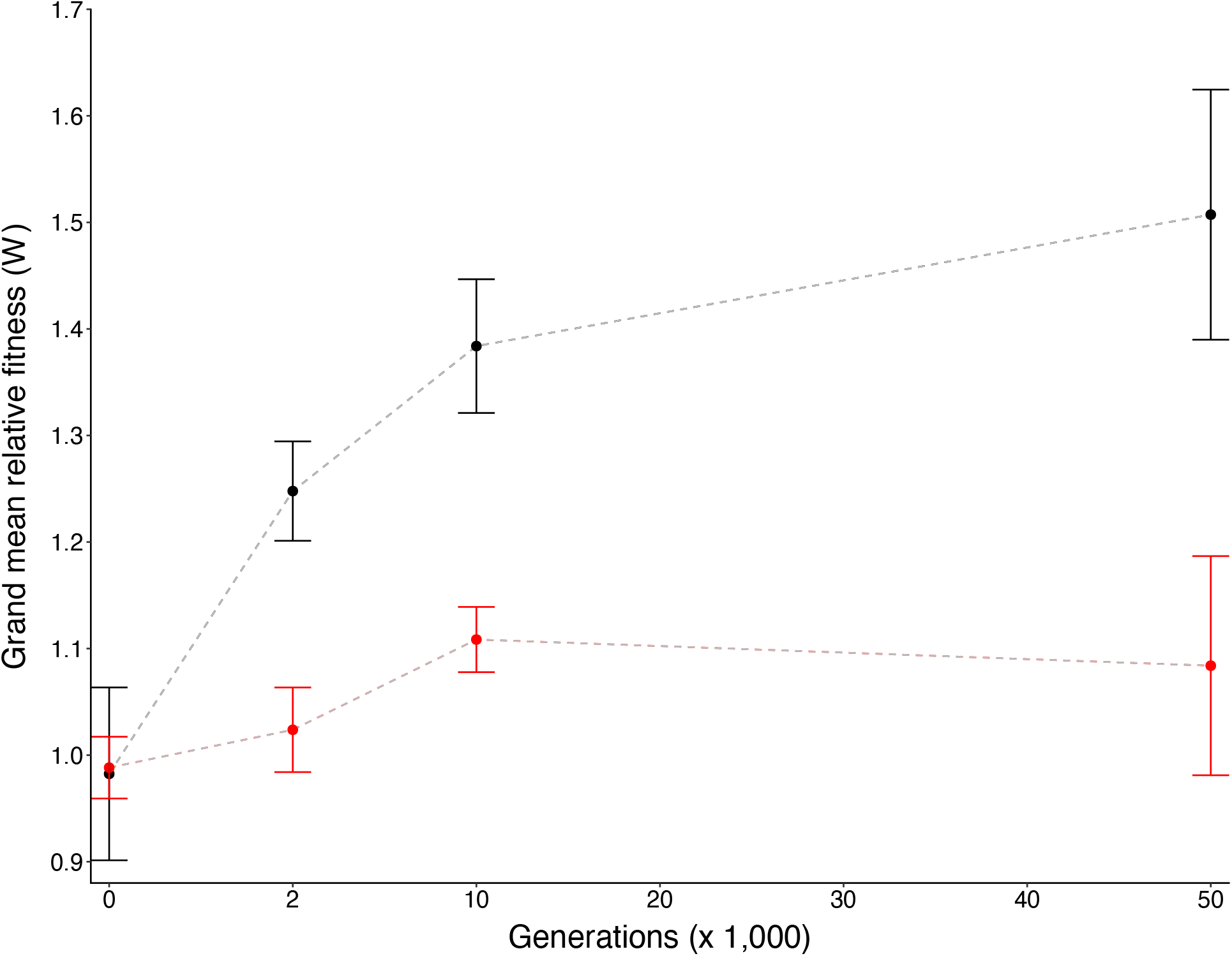
Relative fitness trajectories during 50,000 generations of evolution in an oxic environment. Each point is the grand mean fitness of evolved clones sampled at 2,000, 10,000, and 50,000 generations relative to the LTEE ancestors. Black and red symbols correspond to fitnesses measured in the oxic and anoxic environments, respectively. Error bars show the 95% confidence intervals based on 11, 11, and 9 assayed populations at 2,000, 10,000, and 50,000 generations, respectively.

The fitness of the LTEE populations has increased monotonically in the oxic environment throughout the experiment (Lenski et al. 2015), and our data recapitulate this behavior (fig. 6, supplemental PDF, fig. S3). Although we expected, and confirmed (figs. 5, 6), that fitness relative to the ancestor would be higher in the oxic environment than in the anoxic environment, we did not have a clear expectation for the trajectory of fitness relative to the ancestor in the novel anoxic environment. On the one hand, the anoxic environment shares most aspects of the oxic environment including the limiting resource (glucose), temperature (37°C), and absence of any predators. On the other hand, some aspects of performance might tradeoff between the two conditions. Also, as we saw at the genomic level (figs. 2, 3), anaerobic-specific genes could decay by mutation accumulation, especially in the mutator populations. Given these opposing expectations, one might expect anaerobic fitness to follow a quasi-random walk (Freckleton and Harvey 2006). The fitness trajectories measured in the anoxic environment show some apparent changes in direction, but the differences between consecutive time points are generally within the margin of error (supplemental PDF, fig. S3). In fact, none of the 31 comparisons between sequential points are significant at *p* < 0.05 after performing a Bonferroni correction. In any case, fitness tended to increase even in the anoxic environment in most populations (supplemental PDF, fig. S3). Across all of the clones tested, 23 of 31 had point estimates of their fitness relative to the ancestor in the anoxic environment greater than unity (two-tailed sign test, *p* = 0.0107). However, the grand mean fitness of the evolved bacteria in that environment was significantly greater than unity only at the 10,000-generation time point (fig. 6).

### Heterogeneity of Responses Among Replicate Populations

Each population acquired a unique set of mutations over the course of the LTEE. However, adaptation to the common environment contributed to strong parallelism at the level of genes, especially in the non-mutator populations (Woods et al. 2006; Tenaillon et al. 2016; Good et al. 2017). For example, just 57 genes that make up only ~2% of the coding genome had ~50% of the nonsynonymous mutations that accumulated in the non-mutator populations through 50,000 generations (Tenaillon et al. 2016). The trajectories for fitness also showed strong parallelism (Lenski and Travisano 1994; Wiser et al. 2013; Lenski et al. 2015). For example, the square root of the among-population variance for fitness was only ~5% after 50,000 generations (Lenski et al. 2015), when the grand mean fitness itself had increased by ~70% (Wiser et al. 2013). By contrast, with relaxed selection on anaerobic-specific genes, the accumulation of different sets of mutations should contribute to the populations having greater heterogeneity in fitness when assayed in the anoxic environment than in the oxic environment. We first examined these predictions by performing six one-way ANOVAs (two environments and three time points) to test whether the among-lineage variation was significant (table 1). At 2,000 generations, we saw significant fitness heterogeneity in the anoxic environment, consistent with our expectation. At 10,000 generations, by contrast, there was no significant heterogeneity in either environment. Finally, we saw significant fitness variation in both test environments after 50,000 generations.

**Table 1:**
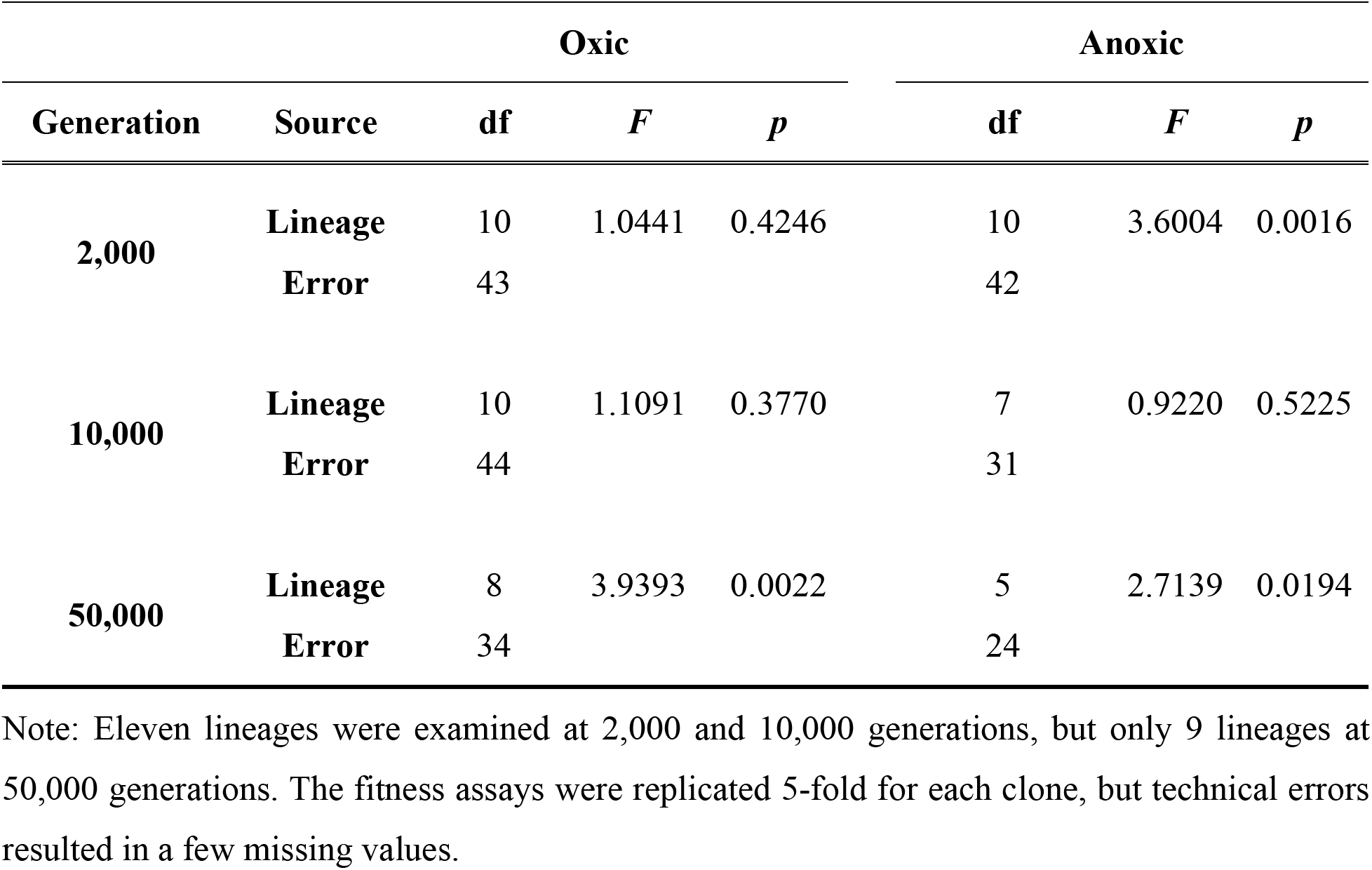
ANOVAs of relative fitness for clones sampled from the LTEE populations at three timepoints and measured in either the oxic or anoxic environment

| Generation | Source | Oxic |  |  | Anoxic |  |  |
| --- | --- | --- | --- | --- | --- | --- | --- |
|  |  | df | *F* | *p* | df | *F* | *p* |
| 2,000 | Lineage | 10 | 1.0441 | 0.4246 | 10 | 3.6004 | 0.0016 |
|  | Error | 43 |  |  | 42 |  |  |
| 10,000 | Lineage | 10 | 1.1091 | 0.3770 | 7 | 0.9220 | 0.5225 |
|  | Error | 44 |  |  | 31 |  |  |
| 50,000 | Lineage | 8 | 3.9393 | 0.0022 | 5 | 2.7139 | 0.0194 |
|  | Error | 34 |  |  | 24 |  |  |

Note: Eleven lineages were examined at 2,000 and 10,000 generations, but only 9 lineages at 50,000 generations. The fitness assays were replicated 5-fold for each clone, but technical errors resulted in a few missing values.

Of course, statistical significance, or the lack thereof, is a crude criterion by which to compare the among-lineage heterogeneity in fitness between the two environments. One can visualize the magnitude of the heterogeneity by estimating the variance attributable to lineages that is greater than expected from the measurement error across replicate assays. However, the statistical uncertainty in estimating variance components is often quite large, and indeed that was the case in our analyses, which showed no significant difference in fitness heterogeneity between the oxic and anoxic environments at any of the generations tested (fig. 7). We also considered the possibility that these analyses were unduly influenced by the subset of populations that evolved hypermutability, which might obscure differences in the among-lineage variation between the two environments. To that end, we repeated the ANOVAs (supplemental PDF, table S2) and the estimation of variance components using only those lineages that did not evolve hypermutability. However, the results of these analyses were not appreciably different (supplemental PDF, fig. S5). In short, contrary to our expectation, we found no compelling evidence of greater among-lineage heterogeneity when fitness was measured in the anoxic than in the oxic environment.

**Figure 7:**
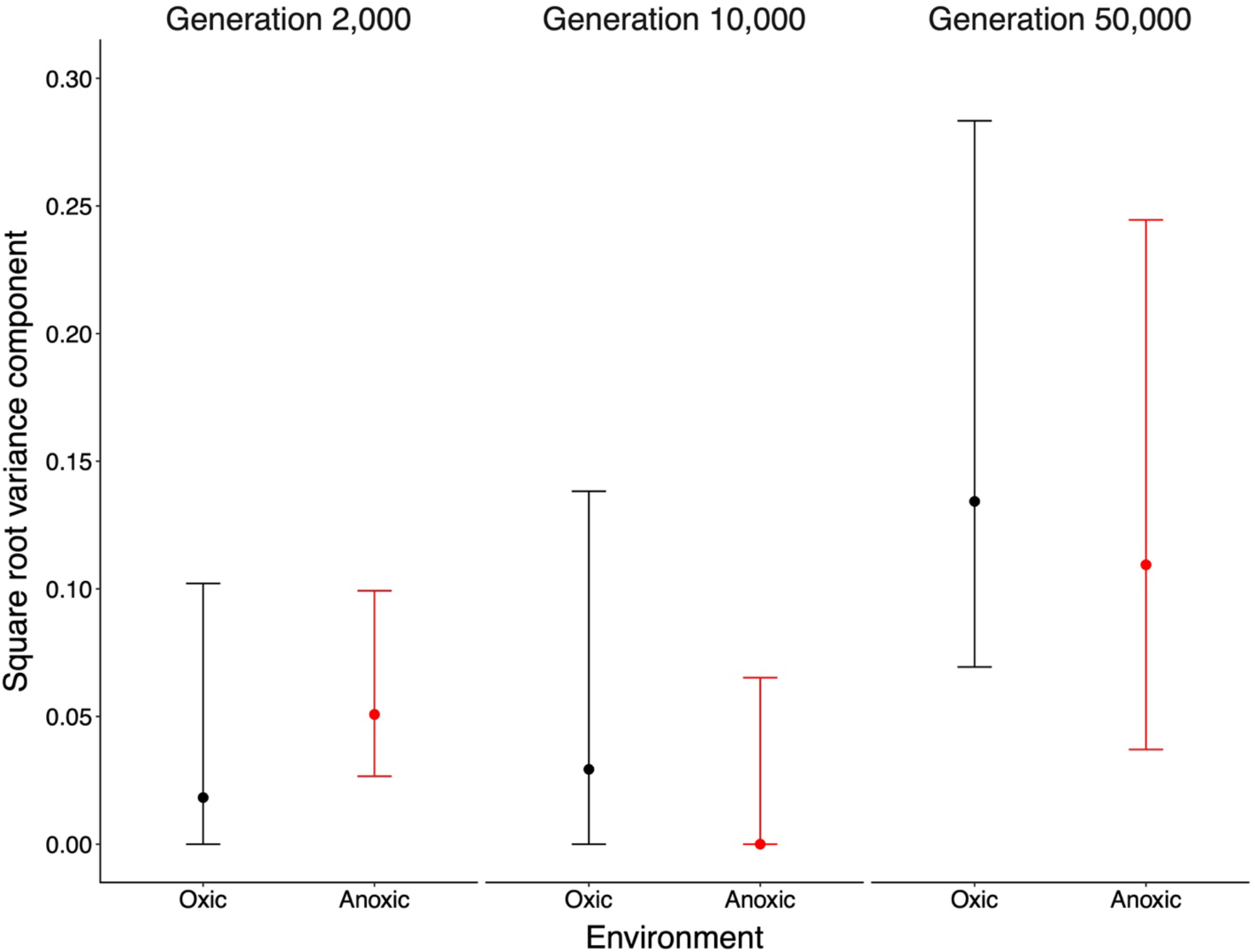
Among-population variance component for fitness in the oxic and anoxic environments. Each point is the square root of the variance component estimated from a corresponding randomeffects ANOVA. Error bars are approximate 95% confidence intervals obtained using the Moriguti-Bulmer procedure (Sokal and Rohlf 1995). Negative values of estimates and confidence limits are truncated at 0, the lower theoretical bound.

## Discussion

If a trait or function is no longer useful to an organism because of a change in its environment, then it may be lost over time. It is unclear, however, what factors determine whether and how quickly traits under relaxed selection will be lost. In this study, we investigated the consequences of relaxed selection for metabolic plasticity by examining the maintenance of anaerobic metabolism—an ancient and core function—using *E. coli* strains that have been evolving in the laboratory under strictly oxic conditions for more than 70,000 generations. On the one hand, one can really imagine that anaerobic metabolism would decay because it has been unused during that time. On the other hand, one can imagine that anaerobic metabolism would be maintained even under relaxed selection owing to the ancient, tightly intertwined physiological and genetic networks that govern aerobic and anaerobic metabolism.

Two mechanisms have been proposed to explain trait loss under relaxed selection (Fong et al. 1995; Cooper and Lenski 2000): antagonistic pleiotropy (AP), and mutational degradation (MD). AP is a selection-driven process whereby improvements to traits that are beneficial in one environment trade off with traits that are useful in other conditions (Williams 1957). An example of AP seen in the LTEE is the loss of the ability to grow on maltose, which has occurred in most populations by mutations in a gene that encodes a transcriptional activator of other genes that encode proteins used to transport and metabolize that sugar (Pelosi et al. 2006; Leiby and Marx 2014). In the absence of maltose, the loss of expression of those genes confers a demonstrable competitive advantage, indicative of AP. A familiar example in the realm of multicellular eukaryotes is senescence, whereby traits that increase reproductive potential early in life reduce survival late in life (Rodríguez et al. 2017). By contrast, MD occurs by a neutral process. In this scenario, traits under relaxed selection accumulate degradative mutations that are neutral in an organism’s current environment, but which reduce the organism’s fitness in environments where those traits are useful. AP and MD are not mutually exclusive, so both can act together to degrade unused functions. In those LTEE populations that evolved greatly elevated mutation rates, one would expect MD to cause much greater decay of unused genes (Cooper and Lenski 2000; Couce et al. 2017). The hypermutable lineages have also increased their fitness in the LTEE environment more than the populations that retained the low ancestral mutation rate, although the difference is small (Wiser et al. 2013; Lenski et al. 2015). Whether the LTEE populations would, on balance, lose fitness under anoxic conditions, and whether those lineages that evolved hypermutability would lose fitness to a greater extent, depends on the form and strength of the regulatory and physiological couplings between aerobic and anaerobic metabolism.

We examined genomic and metagenomic sequence data to detect possible signatures of different modes of selection acting on aerobic- and anaerobic-specific genes. Six of the 12 LTEE populations retained the low ancestral point-mutation rate throughout their history (Tenaillon et al. 2016; Good et al. 2017). All six accumulated many more nonsynonymous mutations in aerobic-specific genes than in anaerobic-specific genes (fig. 2). Moreover, these mutations were concentrated in a subset of genes, with that genetic parallelism indicative of adaptive evolution (Tenaillon et al. 2016; Good et al. 2017). In a number of cases, the inferred benefit of parallel changes was confirmed by measuring the relative fitness of otherwise isogenic strains (Barrick et al. 2009). By contrast, the six populations that evolved hypermutability (Sniegowski et al. 1997; Wielgoss et al. 2013; Tenaillon et al. 2016) accumulated more nonsynonymous mutations in anaerobic-than in aerobic-specific genes (fig. 2). Consistent with purifying selection against mutations in aerobic-specific genes, presumptive knockout mutations (insertions, deletions, nonsense mutations, and structural variants) were underrepresented in those genes in most of the hypermutable populations (fig. 3). More generally, our results agree with several studies that have found a higher frequency of mutations in genes underlying traits under relaxed selection (Shabalina et al. 1997; Cooper and Lenski 2000; Funchain et al. 2000; Maughan et al. 2007; Shewaramani et al. 2017; Cui et al. 2019; Harrison et al. 2019). For example, Maughan et al. (2007) propagated *Bacillus subtilis* vegetatively (i.e., without sporulation) for 6,000 generations. They found that the evolved strains’ inability to generate spores was driven largely by the accumulation of mutations in genes required for sporulation, but not for vegetative growth.

We also identified mutations in *arcA* and *arcB* in almost all of the evolved lines (fig. 4). These genes encode the two-component system responsible for regulating the switch between aerobic and anaerobic metabolism (fig. 1). Previous work found that *arcA* is among the top 15 genes in the LTEE in terms of nonsynonymous substitutions among non-hypermutable lineages, and this parallelism implies positive selection (Tenaillon et al. 2016). Moreover, *arcB* mutations have been implicated in improving growth on acetate—a waste product of glucose metabolism— in some LTEE populations (Plucain et al. 2014; Quandt et al. 2015; Leon et al. 2018).

Given the high rate of mutation accumulation in anaerobic-specific genes in the mutator populations (fig. 2), along with numerous mutations in *arcAB* in both mutator and non-mutator populations (fig. 4), one might reasonably expect that anaerobic metabolism would be degraded, or perhaps even completely lost, in the LTEE populations. However, that was not the case. Our results indicate a more nuanced outcome. On average, the grand mean fitness measured under anoxic conditions tended to increase over the first 10,000 generations of the LTEE, although to a much lesser extent than fitness measured under oxic conditions (fig. 6). Between 10,000 and 50,000 generations, the grand mean fitness measured in the anoxic environment showed no clear trend, even as fitness under the oxic conditions of the LTEE continued to increase, albeit at a slower rate (fig. 6). One might also expect to see greater variation in fitness among the replicate lines when measured in the novel anoxic environment than in the oxic environment where selection was in force during the LTEE. Substantially increased among-population variation in fitness has been reported, for example, when growth substrates (Travisano and Lenski 1996) and temperature were changed (Cooper et al. 2001a). However, we found no meaningful differences in the among-population variance in fitness at any of the time points tested (fig. 7). We also saw no consistent difference in fitness in the anoxic environment between the hypermutable and non-mutator populations (supplemental PDF, fig. S4), despite the compelling evidence for relaxed selection on anaerobic-specific genes in the hypermutable populations (fig. 2). These results broadly suggest that anaerobic metabolism, taken as a whole, did not decay appreciably under relaxed selection, perhaps owing to conserved underlying correlations with traits that contribute to aerobic performance and that experienced a mixture of positive and purifying selection during the LTEE.

Similar correlated responses have been measured for other phenotypic traits in the LTEE populations. For example, the fitness gains made on glucose during the first 2,000 generations led to correlated improvements on lactose (Travisano and Lenski 1996). Leiby and Marx (2014) measured the growth of clones isolated at generations 20,000 and 50,000 on a large array of substrates. They found correlated improvements on some substrates that the populations had not seen for decades, including a few that the ancestors could not even use. Cooper (2002) competed evolved and ancestral strains in four different media, including two with altered glucose concentrations (DM2.5 and DM250), the base medium (DM25) with added bile salts, and a dilute version of the nutritionally complex LB medium. The relative fitness of the evolved lines had, on average, increased in all of these foreign environments. In another study, Meyer et al. (2010) showed that most late-generation LTEE lines had evolved resistance to phage lambda, despite never being exposed to lambda or any other phage during the experiment. They further showed that this positive correlated response was itself associated with a negative correlated response, namely the loss of the capacity to grow on maltose. Lambda uses a maltose transporter to infect cells, and reduced expression of that protein promotes fitness on glucose, compromises growth on maltose, and confers resistance to lambda (Meyer et al. 2010).

What might contribute to the maintenance and even slight improvement of anaerobic performance, despite relaxed selection? The evolved populations showed modest improvement, on average, in the anoxic environment at 10,000 generations (fig. 6). One possibility is that some traits and pathways, such as faster glucose transport and glycolysis, are beneficial in both oxic and anoxic environments. As a consequence, some of the mutated genes that evolved in parallel early in the LTEE might be beneficial in the anoxic environment as well. Only 4 of the 15 genes (*pykF*, *hslU*, *infB*, and *rplF*) with the strongest signatures of parallel evolution (Tenaillon et al. 2016) are in the aerobic-specific gene set, and even some of them might sometimes contribute to anaerobic metabolism. This overlap may explain the correlated improvement in anaerobic fitness early in the LTEE. However, it is unclear whether it is sufficient to explain the maintenance of anaerobic growth capacity after 50,000 generations of relaxed selection, especially in those lines that were hypermutable for much of that time, because knocking out even a single function that is truly required for anaerobic growth should disrupt that metabolic plasticity. In any case, future work might examine the fitness effects of specific mutations that arose in the LTEE, including those that fixed early in the LTEE, on anaerobic fitness to determine if they do, in fact, provide correlated advantages.

Another possible explanation for the maintenance of anaerobic performance hinges on a connection between biochemistry and metabolism. The formulation of the DM25 medium does not provide an alternative terminal electron acceptor that would permit anaerobic respiration. Thus, the bacteria are presumably fermenting glucose in the anoxic environment. Even in the oxic environment of the LTEE, however, the cells might be fermenting rather than respiring glucose, using a process called overflow metabolism (Basan et al. 2015; Swain and Fagan 2019). The use of overflow metabolism would favor the maintenance and even improvement of genes involved in fermentative metabolism. Acetate is a major byproduct of fermentative metabolism, and the fact that several LTEE populations evolved frequency-dependent interactions mediated by crossfeeding on acetate (Elena and Lenski 1997; Rozen et al. 2005; Großkopf et al. 2016; Leon et al. 2018) provides support for this hypothesis. A prediction of this hypothesis is that the bacteria in the LTEE are not respiring, and therefore not using dissolved oxygen, at least during some phase of their population growth. Future studies can test this prediction, and they could also use alternative terminal electron acceptors to measure the respiratory capacity of the evolved lines under anoxic conditions to determine whether their anaerobic growth has become restricted to fermentative metabolism.

A third explanation for the maintenance and even slight improvement in anaerobic fitness of the LTEE populations involves the ArcAB system (fig. 1). These two proteins work in concert to regulate the expression of genes relevant to both aerobic and anaerobic growth (Iuchi and Lin 1988; Gunsalus and Park 1994; Unden and Bongaerts 1997). In particular, ArcA and ArcB together repress the genes that encode TCA-cycle enzymes under anoxic conditions. In their seminal paper describing this system, Iuchi and Lin (1988) isolated *arcA* mutants that produce abnormally high levels of enzymes that are normally repressed under anoxic conditions. These mutants were pleiotropic, so that several aerobic-specific enzymes had increased expression under anoxic conditions including those involved in the TCA cycle and in fatty acid degradation as well as some flavoprotein dehydrogenases and a ubiquinone oxidase. Saxer et al. (2014) also reported extensive metabolic changes caused by *arcA* mutations in short-term evolution experiments with both *E. coli* and *Citrobacter freundii*. They performed proteomic analyses that recapitulated the changes in gene expression reported by Iuchi and Lin (1988), and they also saw increased expression of genes involved in amino-acid metabolism. Saxer et al. (2014) concluded that mutations in global regulators like the ArcAB system could, in one step, expand the niche of an organism by substantially remodeling its cellular metabolism.

All of these potential physiological and genetic explanations for the unexpectedly strong performance of the evolved LTEE bacteria when tested under anoxic conditions might fit under a common theme, which has been called “buttressing pleiotropy” (Lahti et al. 2009). In contrast to the tradeoffs (negative correlations) generated by antagonistic pleiotropy, the idea of buttressing pleiotropy is that “the function of the correlated trait is buttressing or maintaining values of the focal trait” (Lahti et al. 2009). The relevance of buttressing pleiotropy to our results would be strengthened if the genetic architecture underlying aerobic and anaerobic metabolism had many connections. To that end, we used empirical and computationally derived information on proteinprotein interactions to infer and visualize the topology of the aerobic- and anaerobic-specific gene sets in our study (Shannon et al. 2003; Szklarczyk et al. 2017). One immediately sees many connections that demonstrate these two sets are far from independent (fig. 8). It is quite possible, therefore, that selection for improved aerobic growth would buttress anaerobic performance.

**Figure 8:**
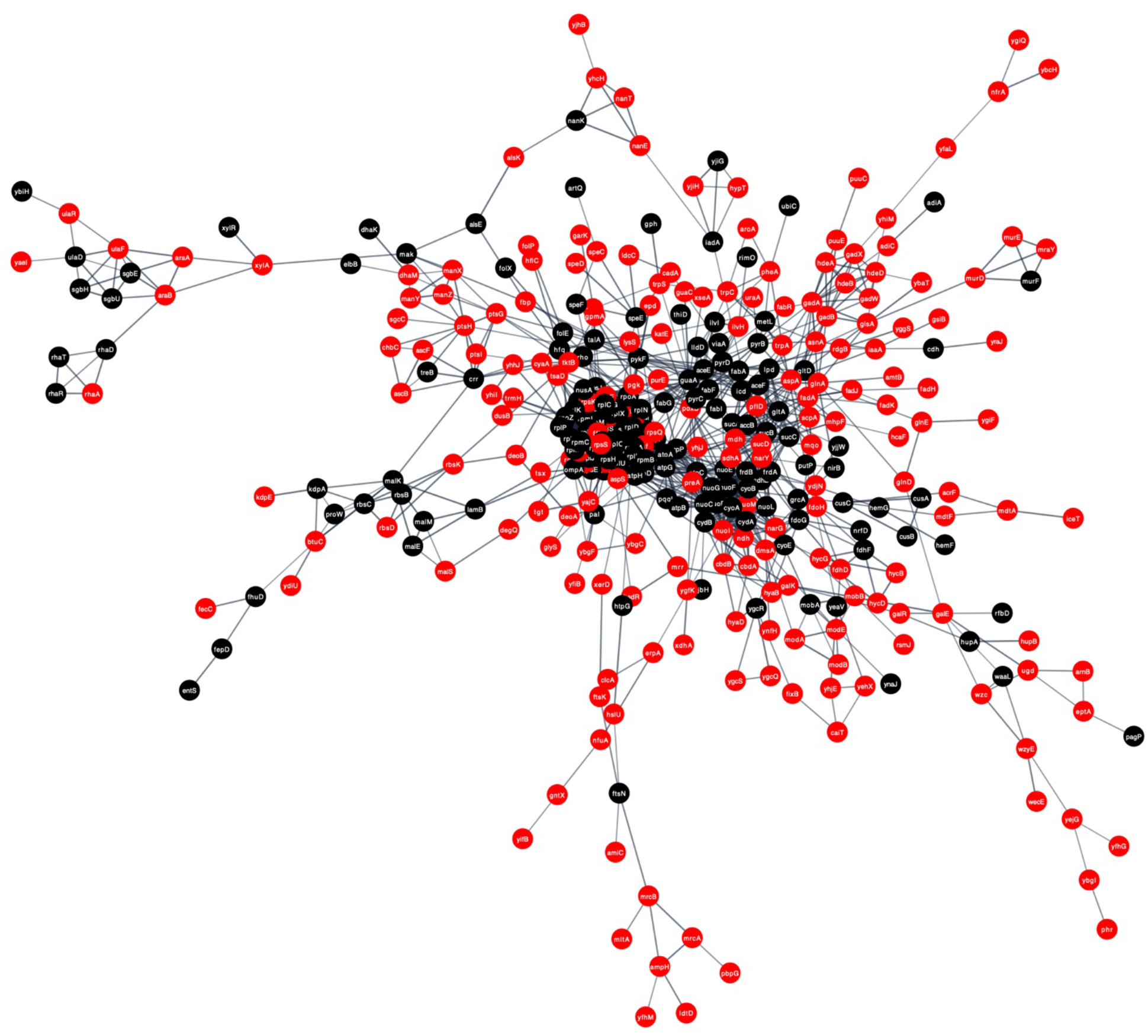
Network topology of aerobic- and anaerobic-specific genes. We used Cytoscape (Shannon et al. 2003) to visualize protein-protein interactions between aerobic-(black) and anerobic-specific (red) genes predicted by STRING (Szklarczyk et al. 2017). Each edge reflects a confidence score of at least 0.7 (i.e., high) that an interaction exists, obtained using a maximum likelihood approach and seven lines of evidence. In the supplemental PDF, Fig. S6 shows additional genes that are not integrated within the main network shown here.

A related idea, also relevant to our system, is robustness. *E. coli* is often described as a facultative anaerobe because it can grow not only in the well-oxygenated conditions widely used in most laboratories, but also when deprived of oxygen in special growth chambers. However, *E. coli* might be better described as a facultative aerobe, because its natural home is the anoxic mammalian colon. To be sure, many *E. coli* cells periodically exit their hosts, and the ability to survive in the presence of oxygen is essential for colonizing new hosts. However, we would suggest that most of the 100-million year or so history of this species has been spent living under anerobic conditions—even if there are as many *E. coli* cells outside as inside mammalian hosts at any given time, those that are outside are much more likely to be evolutionary dead-ends. If so, then evolution might have favored a more robust anaerobic metabolism, one that could not easily be disrupted by short-sighted selection for improved fitness in the oxic environment. A test of this anaerobic-robustness hypothesis would be to perform an experiment identical to the LTEE, except in a strictly anoxic environment. If populations that evolved for 50,000 generations in the absence of oxygen lost the ability to grow in its presence, then that would indicate that aerobic metabolism is less robust that anaerobic metabolism.

In a different vein, it is known that some enzymes exhibit substrate promiscuity (Nam et al. 2012). Changes to global regulators—including the ArcAB system and others that have changed during the LTEE (Cooper et al. 2003, 2008; Philippe et al. 2007; Crozat et al. 2011)—might increase expression of some promiscuous enzymes, and those latent functions might yield benefits in multiple environments. Future work might compare the transcriptomes of ancestral and evolved bacteria in oxic and anoxic environments, as well as examine the effects of specific mutations to ArcAB and other global regulators on the transcriptomes. Another area for future work would examine the speed with which the ancestral and evolved lines can respond to sudden changes in the environment from oxic to anoxic and vice versa. Yet another interesting direction might use a combination of genetic engineering and experimental evolution to generate obligate aerobic and obligate anaerobic strains, which could then be studied to better understand the constraints on each type of metabolism.

All in all, our results show that anaerobic metabolism was surprisingly robust during 50,000 generations of adaptation to a strictly oxic environment. The capacity for anaerobic growth persisted even in those lineages that evolved hypermutability, despite genetic signatures that showed increased mutation accumulation in anaerobic-specific genes in those lines. Thus, relaxed selection on a functional trait does not always result in its loss. We suggest that one must know how the genes and proteins responsible for traits that experience relaxed selection interact with the rest of an organism’s genome and physiology, in order to understand the potential for loss, maintenance, or even correlated improvement of such traits. More generally, investigating the role of genetic architecture in the evolutionary process might help us better predict how organisms will respond to novel selection pressures and environmental perturbations, including climate change. Such understanding may also suggest new ways of promoting or constraining the evolutionary trajectories of organisms and their traits, which could be used for synthetic biology, on the one hand, and to limit the evolution of pathogens, on the other hand.

## Acknowledgments

We thank Terence Marsh, Charles Ofria, Gemma Reguera, and Chris Waters for feedback as this research progressed; Zachary Blount for helpful comments on the manuscript; and members of the Lenski lab for valuable discussions. We thank Terence Marsh for providing access to an anaerobic chamber, and the MSU Department of Microbiology and Molecular Genetics for related supplies. This work was supported in part by a grant from the National Science Foundation (currently DEB-1951307), the BEACON Center for the Study of Evolution in Action (DBI-0939454), and the USDA National Institute of Food and Agriculture (MICL02253). Any opinions, findings, and conclusions or recommendations expressed in this material are those of the authors and do not necessarily reflect the views of the funders.

## Supplemental information for the paper

**Figure S1:**
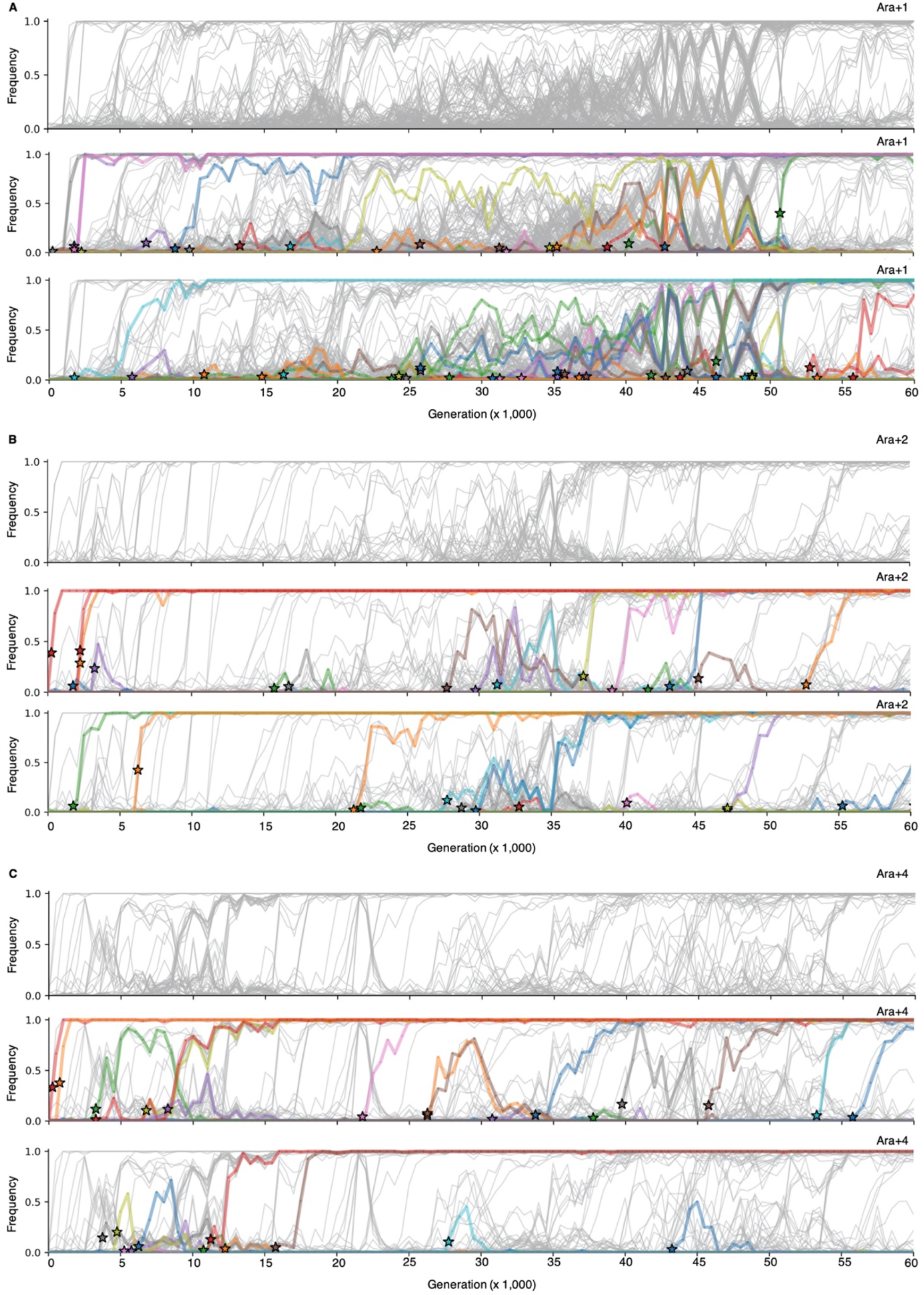
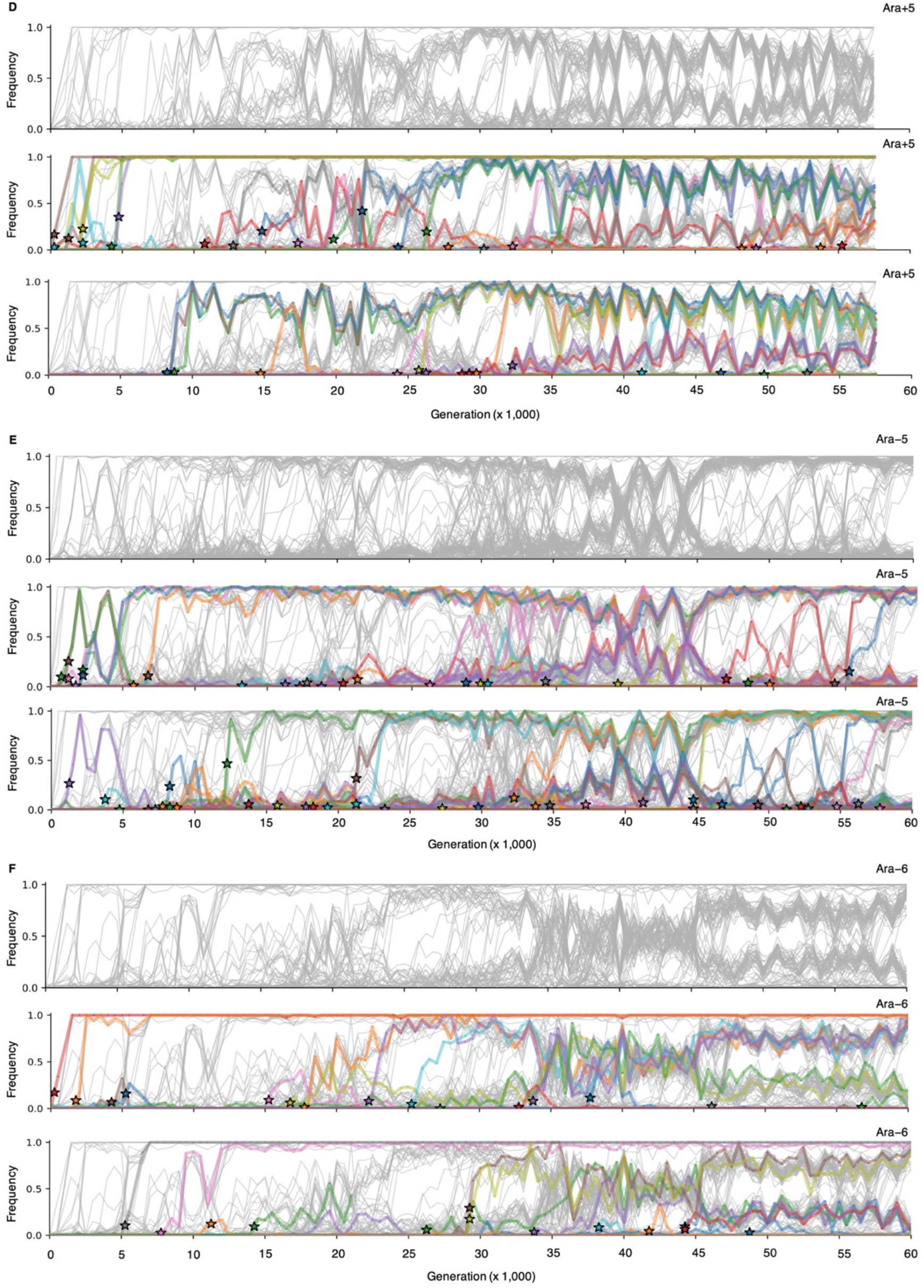
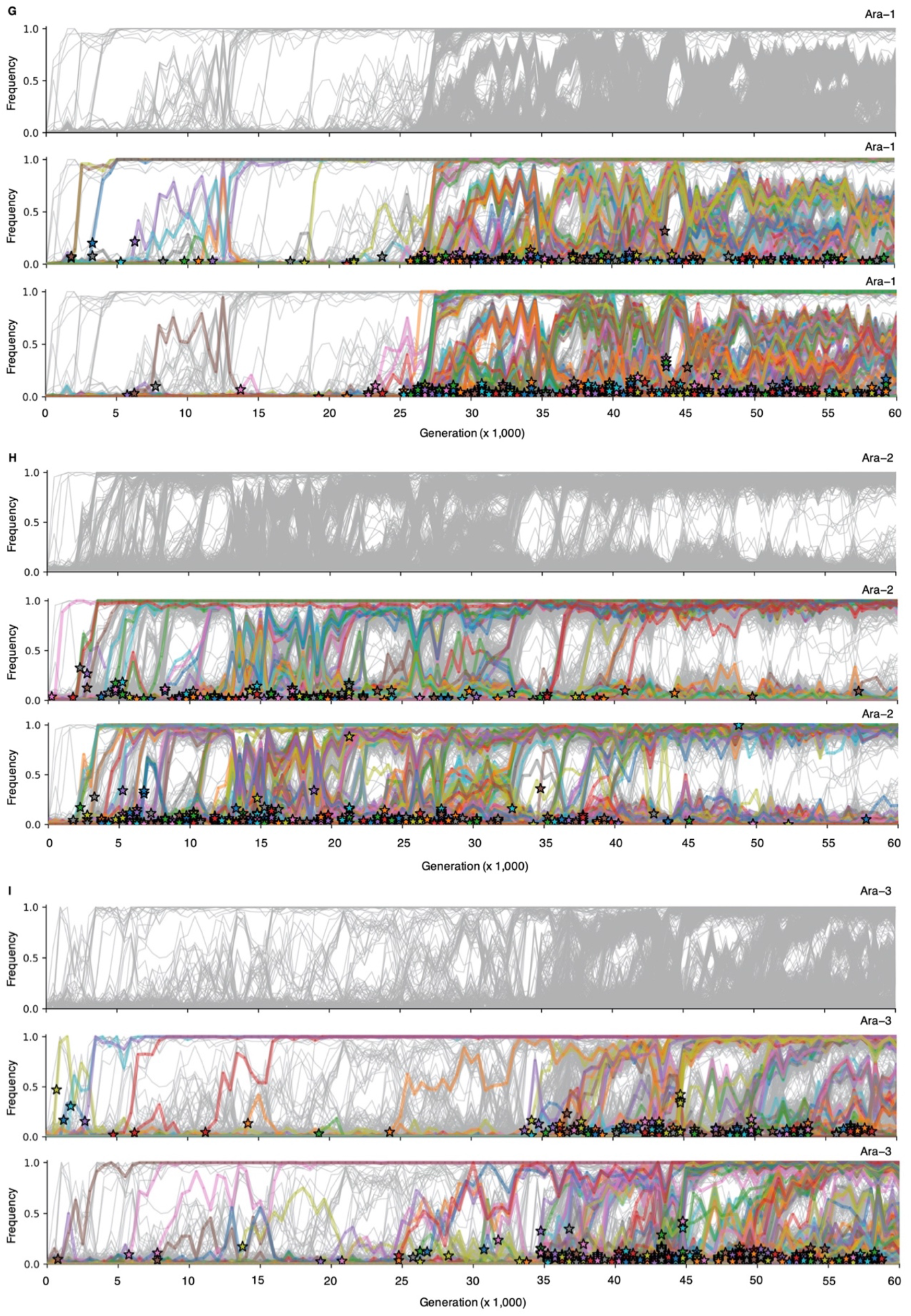
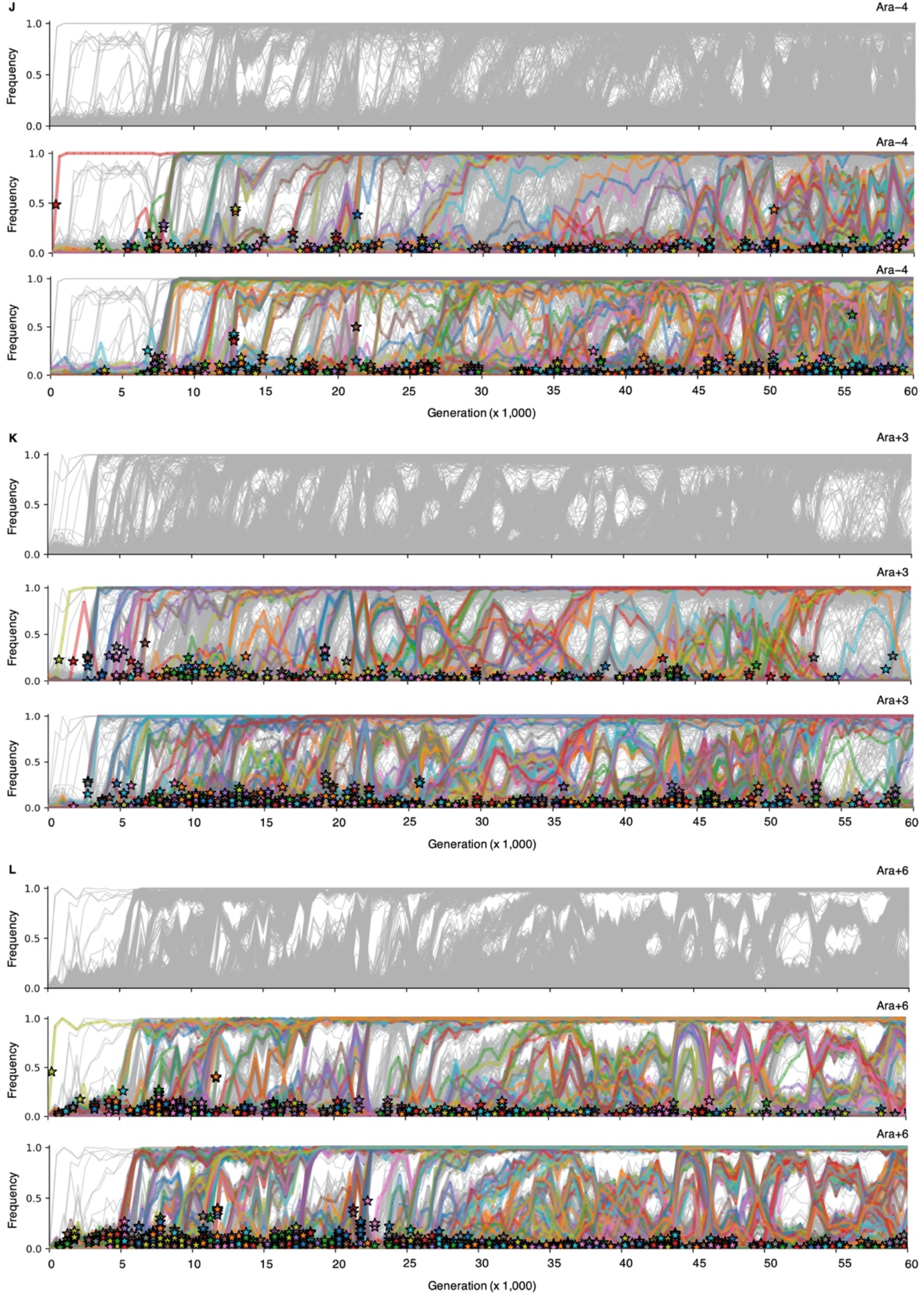
Evolutionary dynamics of mutations in aerobic-specific genes. Each panel (A–L) shows the allele-frequency trajectories for new mutations that arose in the indicated LTEE population and reached an observable frequency. Panels A–F show the six populations that never evolved hypermutability; panels G–L show the six populations that became mutators at various time points. In each panel, the top sub-panel shows in gray all observed mutations; the middle subpanel colors only those mutations in aerobic-specific genes; and the bottom sub-panel colors only those mutations in anaerobic-specific genes. The underlying metagenomic data and statistical criteria are from Good et al. (2017). The stars in the middle and bottom sub-panels mark the “appearance time” of mutations, which Good et al. (2017) defined as 250 generations before the first sample in which a given mutation reached observable frequency (i.e., the midpoint between that sample and the preceding sample, given 500 generations between successive samples).

**Figure S2:**
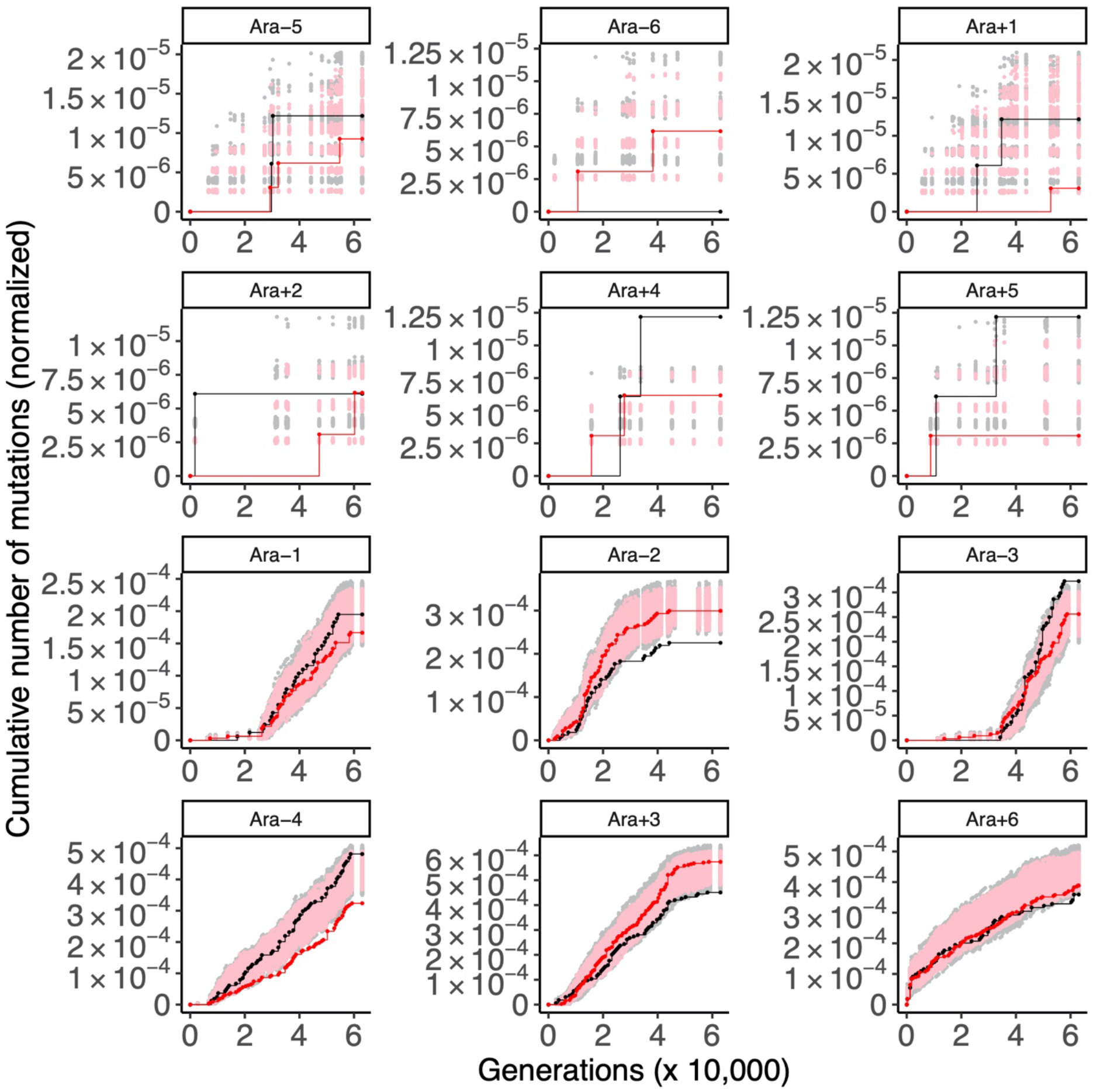
Cumulative number of mutations in aerobic- and anaerobic-specific genes in the LTEE whole-population samples. Each panel shows the number of synonymous mutations in aerobic-(black) and anaerobic-specific (red) genes in the indicated population. For comparison, random sets of genes of equal cardinality to the aerobic-(227) or anaerobic-specific (345) gene sets were sampled 1,000 times, and the cumulative number of synonymous mutations was calculated to generate a null distribution of the expected number of mutations for each population. The gray and pink points show 95% of these null distributions (excluding 2.5% in each tail) for aerobic and anaerobic comparisons, respectively.

**Figure S3:**
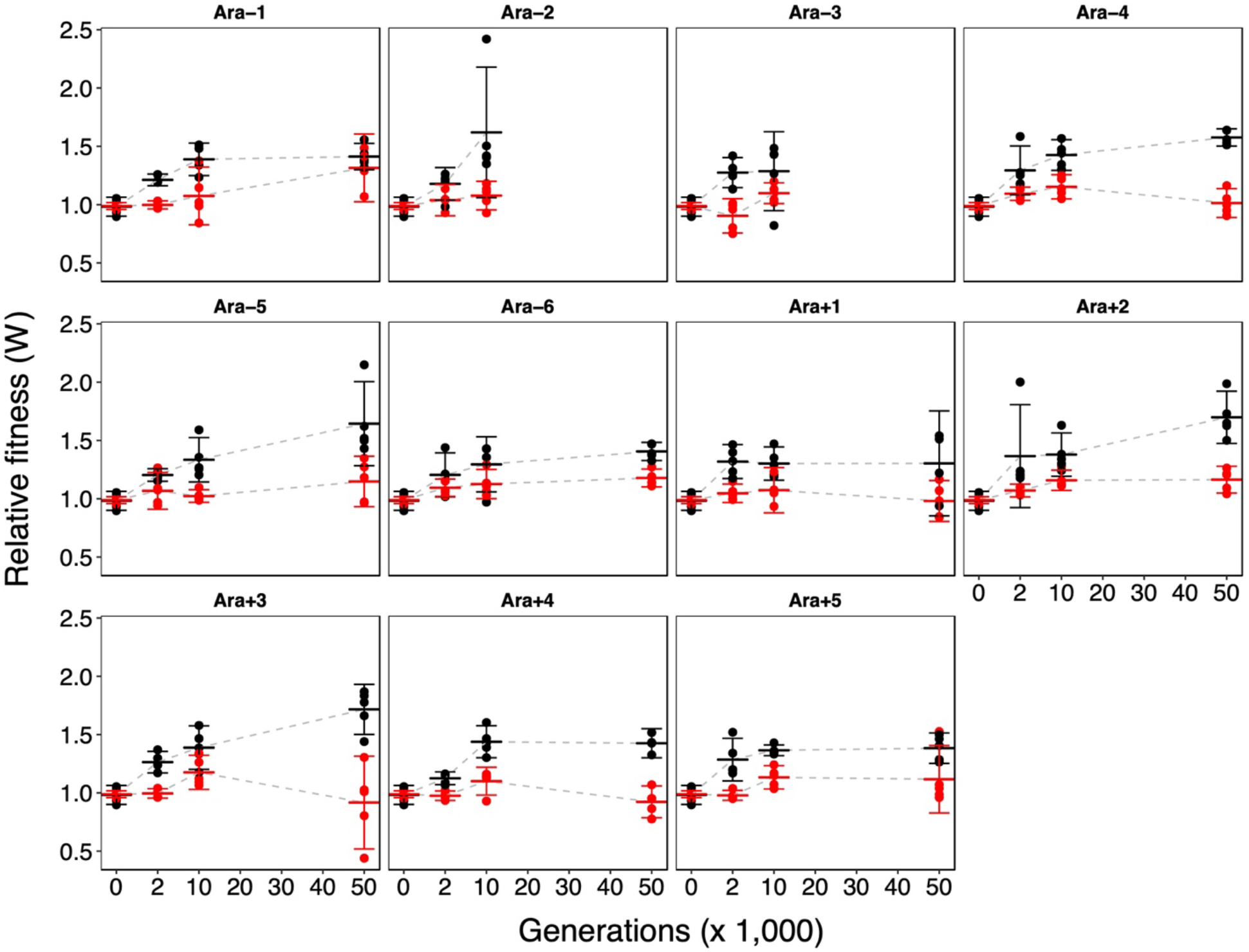
Relative fitness trajectories of individual LTEE populations in oxic and anoxic environments. Black and red points show competitions performed in oxic and anoxic environments, respectively. Each point is a replicate competition assay in which a clone from the indicated population was competed against the reciprocally marked ancestral strain. Two trajectories are truncated owing to technical difficulties (see Materials and Methods). Wide hash marks are means; error bars are 95% confidence intervals.

**Figure S4:**
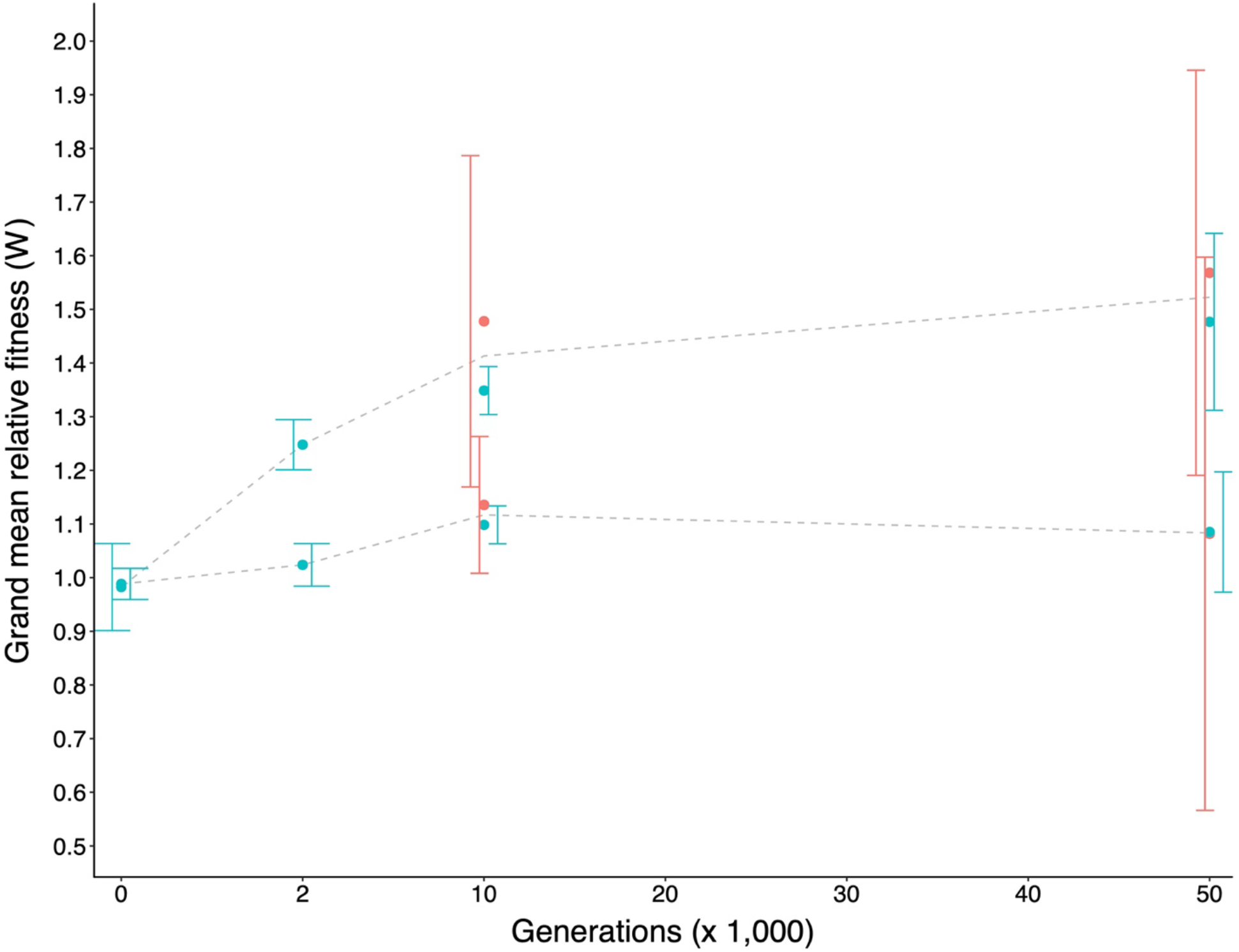
Relative fitness trajectories for the LTEE populations shown by their mutator status. Each point is the grand-mean fitness value of a set of evolved clones sampled at 2,000, 10,000, and 50,000 generations relative to the ancestors. Non-mutators and mutators are shown in teal and orange, respectively. The upper and lower trajectories were measured in the oxic and anoxic environments, respectively. Error bars are 95% confidence intervals.

**Figure S5:**
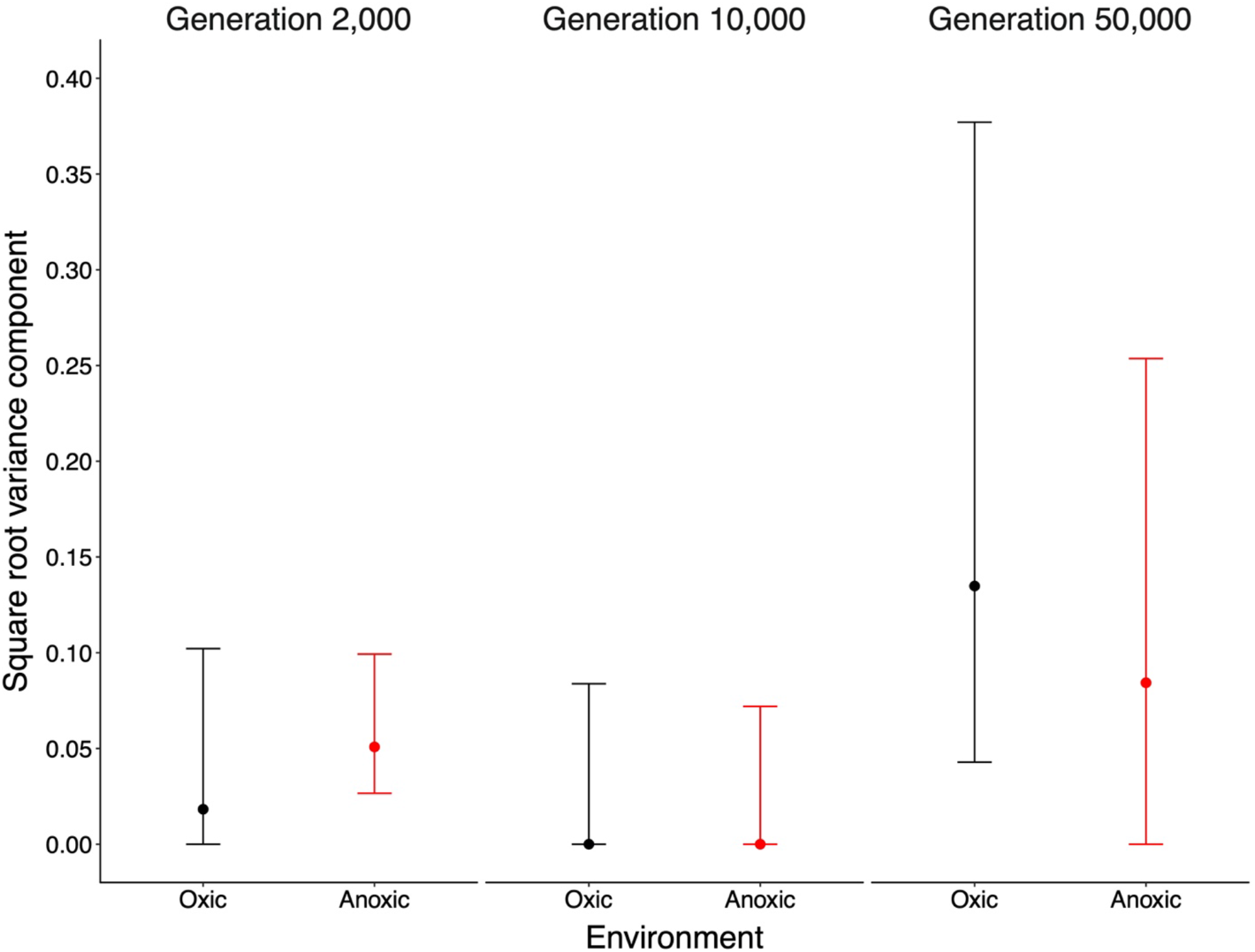
Among-population variance component for fitness, excluding mutator populations. See Figure 7 for additional details. The among-population variance is similar in the two environments whether the mutator populations are included (fig. 7) or excluded (this figure).

**Figure S6:**
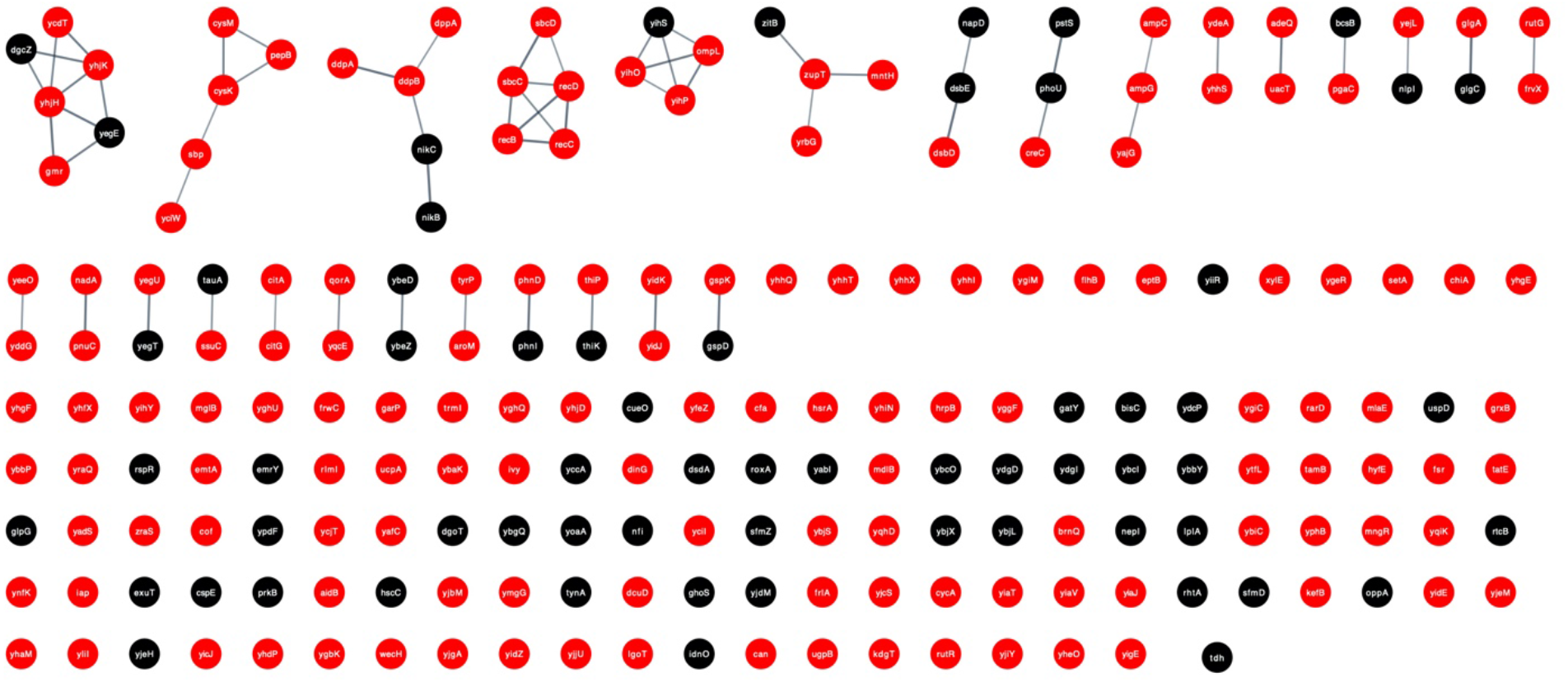
Network topology aerobic-(black) and anerobic-specific (red) genes that are not connected to the large core network shown in Figure 8. See the legend to that figure for additional details.

**Table S1:** List of *E. coli* clones used in study

| Clone ID | Population | Generation |
| --- | --- | --- |
| REL606 | Ara<sup>-</sup> ancestor | 0 |
| REL607 | Ara<sup>+</sup> ancestor | 0 |
| REL1158A | Ara+1 | 2,000 |
| REL1159A | Ara+2 | 2,000 |
| REL1160A | Ara+3 | 2,000 |
| REL1161A | Ara+4 | 2,000 |
| REL1162A | Ara+5 | 2,000 |
| REL1163A | Ara+6 | 2,000 |
| REL1164A | Ara-1 | 2,000 |
| REL1165A | Ara-2 | 2,000 |
| REL1166A | Ara-3 | 2,000 |
| REL1167A | Ara-4 | 2,000 |
| REL1168A | Ara-5 | 2,000 |
| REL1169A | Ara-6 | 2,000 |
| REL4530A | Ara+1 | 10,000 |
| REL4531A | Ara+2 | 10,000 |
| REL4532A | Ara+3 | 10,000 |
| REL4533A | Ara+4 | 10,000 |
| REL4534A | Ara+5 | 10,000 |
| REL4535A | Ara+6 | 10,000 |
| REL4536A | Ara-1 | 10,000 |
| REL4537A | Ara-2 | 10,000 |
| REL4538A | Ara-3 | 10,000 |
| REL4539A | Ara-4 | 10,000 |
| REL4540A | Ara-5 | 10,000 |
| REL4541A | Ara-6 | 10,000 |
| REL11392 | Ara+1 | 50,000 |
| REL11342 | Ara+2 | 50,000 |
| REL11345 | Ara+3 | 50,000 |
| REL11348 | Ara+4 | 50,000 |
| REL11367 | Ara+5 | 50,000 |
| REL11370 | Ara+6 | 50,000 |
| REL11330 | Ara-1 | 50,000 |
| REL11333 | Ara-2 | 50,000 |
| REL11364 | Ara-3 | 50,000 |
| REL11336 | Ara-4 | 50,000 |
| REL11339 | Ara-5 | 50,000 |
| REL11389 | Ara-6 | 50,000 |

**Table S2:** ANOVAs of relative fitness for clones sampled from non-mutator populations at three timepoints and measured in the oxic or anoxic environment

| **Generation** | **Source** | **Oxic** |  |  | **Anoxic** |  |  |
| --- | --- | --- | --- | --- | --- | --- | --- |
|  |  | **df** | ***F*** | ***p*** | **df** | ***F*** | ***p*** |
| **2,000** | **Lineage** | 10 | 1.0441 | 0.4246 | 10 | 3.6004 | 0.0016 |
|  | **Error** | 43 |  |  | 42 |  |  |
| **10,000** | **Lineage** | 7 | 0.5832 | 0.7644 | 7 | 0.7684 | 0.6180 |
|  | **Error** | 32 |  |  | 31 |  |  |
| **50,000** | **Lineage** | 5 | 3.2700 | 0.0234 | 5 | 2.6544 | 0.0478 |
|  | **Error** | 22 |  |  | 24 |  |  |

Note: Data were obtained for 11, 8, and 6 lineages at 2,000, 10,000, and 50,000 generations, respectively. See Table 1 for additional details.

## Literature Cited

Altenhoff, A. M., N. M. Glover, C. M. Train, K. Kaleb, A. Warwick Vesztrocy, D. Dylus, T. M. De Farias, et al. 2018. The OMA orthology database in 2018: Retrieving evolutionary relationships among all domains of life through richer web and programmatic interfaces. Nucleic Acids Research 46(D1):D477–D485.

Barrick, J. E., D. Yu, S. Yoon, H. Jeong, T. Oh, D. Schneider, R. E. Lenski, et al. 2009. Genome evolution and adaptation in a long-term experiment with *Escherichia coli*. Nature 461:1243–1247.

Basan, M., S. Hui, H. Okano, Z. Zhang, Y. Shen, J. R. Williamson, and T. Hwa. 2015. Overflow metabolism in *Escherichia coli* results from efficient proteome allocation. Nature 528:99–104.

Bennett, A. F., K. M. Dao, and R. E. Lenski. 1990. Rapid evolution in response to high-temperature selection. Nature 346:79–81.

Blazejewski, T., H. I. Ho, and H. H. Wang. 2019. Synthetic sequence entanglement augments stability and containment of genetic information in cells. Science 365:595–598.

Cooper, T. F., D. E. Rozen, and R. E. Lenski. 2003. Parallel changes in gene expression after 20,000 generations of evolution in *Escherichia coli*. Proceedings of the National Academy of Sciences of the USA 100:1072–1077.

Cooper, T. F., S. K. Remold, R. E. Lenski, and D. Schneider. 2008. Expression profiles reveal parallel evolution of epistatic interactions involving the CRP regulon in *Escherichia coli*. PLoS Genetics 4:e35.

Cooper, V. S. 2002. Long-term experimental evolution in *Escherichia coli*. X. Quantifying the fundamental and realized niche. BMC Evolutionary Biology 2:12.

Cooper, V. S., and R. E. Lenski. 2000. The population genetics of ecological specialization in evolving *Escherichia coli* populations. Nature 407:736–739.

Cooper, V. S., A. F. Bennett, and R. E. Lenski. 2001a. Evolution of thermal dependence of growth rate of *Escherichia coli* populations during 20,000 generations in a constant environment. Evolution 55:889–896.

Cooper, V. S., D. Schneider, M. Blot, and R. E. Lenski. 2001b. Mechanisms causing rapid and parallel losses of ribose catabolism in evolving populations of *Escherichia coli* B. Journal of Bacteriology 183:2834–2841.

Couce, A., L. V. Caudwell, C. Feinauer, T. Hindré, J. P. Feugeas, M. Weigt, R. E. Lenski, et al. 2017. Mutator genomes decay, despite sustained fitness gains, in a long-term experiment with bacteria. Proceedings of the National Academy of Sciences of the USA 114:E9026–E9035.

Crozat, E., T. Hindré, L. Kühn, J. Garin, R. E. Lenski, and D. Schneider. 2011. Altered regulation of the OmpF porin by Fis in *Escherichia coli* during an evolution experiment and between B and K-12 strains. Journal of Bacteriology 193:429–440.

Cui, R., T. Medeiros, D. Willemsen, L. N. M. Iasi, G. E. Collier, M. Graef, M. Reichard, et al. 2019. Relaxed selection limits lifespan by increasing mutation load. Cell 178:385–399.

Darwin, C. 1859. On the Origin of Species. John Murray, London.

Elena, S. F., and R. E. Lenski. 1997. Long-term experimental evolution in *Escherichia coli*. VII. mechanisms maintaining genetic variability within populations. Evolution 51:1058–1067.

Elena, S. F., and R. E. Lenski. 2003. Evolution experiments with microorganisms: the dynamics and genetic bases of adaptation. Nature Reviews Genetics 4:457–469.

Fong, D. W., T. C. Kane, and D. C. Culver. 1995. Vestigialization and loss of nonfunctional characters. Annual Review of Ecology and Systematics 26:249–268.

Frankel, N., G. K. Davis, D. Vargas, S. Wang, F. Payre, and D. L. Stern. 2010. Phenotypic robustness conferred by apparently redundant transcriptional enhancers. Nature 466:490–493.

Fraser, H. B., and E. E. Schadt. 2010. The quantitative genetics of phenotypic robustness. PLoS ONE 5:e8635.

Freckleton, R. P., and P. H. Harvey. 2006. Detecting non-Brownian trait evolution in adaptive radiations. PLoS Biology 4:2104–2111.

Funchain, P., A. Yeung, J. L. Stewart, R. Lin, M. M. Slupska, and J. H. Miller. 2000. The consequences of growth of a mutator strain of *Escherichia coli* as measured by loss of function among multiple gene targets and loss of fitness. Genetics 154:959–970.

Geng, P., S. P. Leonard, D. M. Mishler, and J. E. Barrick. 2019. Synthetic genome defenses against selfish DNA elements stabilize engineered bacteria against evolutionary failure. ACS Synthetic Biology 8:521–531.

Good, B. H., M. J. McDonald, J. E. Barrick, R. E. Lenski, and M. M. Desai. 2017. The dynamics of molecular evolution over 60,000 generations. Nature 551:45–50.

Gordon, D. M., and A. Cowling. 2003. The distribution and genetic structure of *Escherichia coli* in Australian vertebrates: Host and geographic effects. Microbiology 149:3575–3586.

Großkopf, T., J. Consuegra, J. Gaffé, J. C. Willison, R. E. Lenski, O. S. Soyer, and D. Schneider. 2016. Metabolic modelling in a dynamic evolutionary framework predicts adaptive diversification of bacteria in a long-term evolution experiment. BMC Evolutionary Biology 16:163.

Gunsalus, R. P., and S. J. Park. 1994. Aerobic-anaerobic gene regulation in *Escherichia coli*: Control by the ArcAB and Fnr regulons. Research in Microbiology 145:437–450.

Harrison, M. C., E. B. Mallon, D. Twell, and R. L. Hammond. 2019. Deleterious mutation accumulation in *Arabidopsis thaliana* pollen genes: A role for a recent relaxation of selection. Genome Biology and Evolution 11:1939–1951.

Iuchi, S., and E. C. Lin. 1988. *arcA* (dye), a global regulatory gene in *Escherichia coli* mediating repression of enzymes in aerobic pathways. Proceedings of the National Academy of Sciences of the USA 85:1888–1892.

Jeong, H., V. Barbe, C. H. Lee, D. Vallenet, D. S. Yu, S.-H. Choi, A. Couloux, et al. 2009. Genome sequences of *Escherichia coli* B strains REL606 and BL21(DE3). Journal of Molecular Biology 394:644–652.

Jia, D., M. Lu, K. H. Jung, J. H. Park, L. Yu, J. N. Onuchic, B. A. Kaipparettu, et al. 2019. Elucidating cancer metabolic plasticity by coupling gene regulation with metabolic pathways. Proceedings of the National Academy of Sciences of the USA 116:3909–3918.

Kang, Y., K. D. Weber, Y. Qiu, P. J. Kiley, and F. R. Blattner. 2005. Genome-wide expression analysis indicates that FNR *of Escherichia coli* K-12 regulates a large number of genes of unknown function. Journal of Bacteriology 187:1135–1160.

Keseler, I. M., A. Mackie, A. Santos-Zavaleta, R. Billington, C. Bonavides-Martínez, R. Caspi, C. Fulcher, et al. 2017. The EcoCyc database: Reflecting new knowledge about *Escherichia coli* K-12. Nucleic Acids Research 45:D543–D550.

Kwon, O., D. Georgellis, E.C. Lin. 2003. Rotational on-off switching of a hybrid membrane sensor kinase Tar-ArcB in *Escherichia coli*. Journal of Biological Chemistry 278:13192–13195.

Lahti, D. C., N. A. Johnson, B. C. Ajie, S. P. Otto, A. P. Hendry, D. T. Blumstein, R. G. Coss, et al. 2009. Relaxed selection in the wild. Trends in Ecology and Evolution 24:487–496.

Lamrabet, O., M. Martin, R. E. Lenski, and D. Schneider. 2019. Changes in intrinsic antibiotic susceptibility during a long-term evolution experiment with *Escherichia coli*. mBio 10:e00189–19.

Leiby, N., and C. J. Marx. 2014. Metabolic erosion primarily through mutation accumulation, and not tradeoffs, drives limited evolution of substrate specificity in *Escherichia coli*. PLoS Biology 12:e1001789.

Lenski, R. E. 2017. Experimental evolution and the dynamics of adaptation and genome evolution in microbial populations. ISME Journal 11:2181–2194.

Lenski, R. E., and M. Travisano. 1994. Dynamics of adaptation and diversification: a 10,000-generation experiment with bacterial populations. Proceedings of the National Academy of Sciences of the USA 91:6808–6814.

Lenski, R. E., M. R. Rose, S. C. Simpson, and S. C. Tadler. 1991. Long-term experimental evolution in *Escherichia coli*. I. Adaptation and divergence during 2,000 generations. American Naturalist 138:1315–1341.

Lenski, R. E., C. Ofria, R. T. Pennock, and C. Adami. 2003. The evolutionary origin of complex features. Nature 423:139–144.

Lenski, R. E., J. E. Barrick, and C. Ofria. 2006. Balancing robustness and evolvability. PLoS Biology 4:e428.

Lenski, R. E., M. J. Wiser, N. Ribeck, Z. D. Blount, J. R. Nahum, J. J. Morris, L. Zaman, et al. 2015. Sustained fitness gains and variability in fitness trajectories in the long-term evolution experiment with *Escherichia coli*. Proceedings of the Royal Society B 282:20152292.

Leon, D., S. D’Alton, E. M. Quandt, and J. E. Barrick. 2018. Innovation in an *E. coli* evolution experiment is contingent on maintaining adaptive potential until competition subsides. PLoS Genetics 14:e1007348.

Maddamsetti, R., R. E. Lenski, and J. E. Barrick. 2015. Adaptation, clonal interference, and frequency-dependent interations in a long-term evolution experiment with *Escherichia coli*. Genetics 200:619–631.

Maddamsetti, R., P. J. Hatcher, A. G. Green, B. L. Williams, D. S. Marks, and R. E. Lenski. 2017. Core genes evolve rapidly in the long-term evolution experiment with *Escherichia coli*. Genome Biology and Evolution 9:1072–1083.

Maughan, H., J. Masel, C. W. Birky, and W. L. Nicholson. 2007. The roles of mutation accumulation and selection in loss of sporulation in experimental populations of *Bacillus subtilis*. Genetics 177: 937–948.

Maughan, H., C. W. Birky, and W. L. Nicholson. 2009. Transcriptome divergence and the loss of plasticity in *Bacillus subtilis* after 6,000 generations of evolution under relaxed selection for sporulation. Journal of Bacteriology 191:428–433.

Melville, S. B., and R. P. Gunsalus. 1990. Mutations in fnr that alter anaerobic regulation of electron transport-associated genes in *Escherichia coli*. Journal of Biological Chemistry 265:18733–18736.

Meyer, J. R., A. A. Agrawal, R. T. Quick, D. T. Dobias, D. Schneider, and R. E. Lenski. 2010. Parallel changes in host resistance to viral infection during 45,000 generations of relaxed selection. Evolution 64:3024–3034.

Müller, H. E. 1977. Age and evolution of bacteria. Experientia 33:979–984.

Nam, H., N. E. Lewis, J. A. Lerman, D.-H. Lee, R. L. Chang, D. Kim, and B. O. Palsson. 2012. Network context and selection in the evolution to enzyme specificity. Science 337:1101–1104.

Nijhout, F. H., F. Sadre-Marandi, J. Best, and M. C. Reed. 2017. Systems biology of phenotypic robustness and plasticity. Integrative and Comparative Biology 57:171–184.

Ostrowski, E. A., C. Ofria, and R. E. Lenski. 2015. Genetically integrated traits and rugged adaptive landscapes in digital organisms. BMC Evolutionary Biology 15:83.

Paudel, B. B., and V. Quaranta. 2019. Metabolic plasticity meets gene regulation. Proceedings of the National Academy of Sciences of the USA 116:3370–3372.

Payen, V. L., P. E. Porporato, B. Baselet, and P. Sonveaux. 2016. Metabolic changes associated with tumor metastasis, part 1: Tumor pH, glycolysis and the pentose phosphate pathway. Cellular and Molecular Life Sciences 73:1333–1348.

Pelosi, L., L. Kühn, D. Guetta, J. Garin, J. Geiselmann, R. E. Lenski, and D. Schneider. 2006. Parallel changes in global protein profiles during long-term experimental evolution in *Escherichia coli*. Genetics 173:1851–1869.

Philippe, N., E. Crozat, R. E. Lenski, and D. Schneider. 2007. Evolution of global regulatory networks during a long-term experiment with *Escherichia coli*. BioEssays 29:846–860.

Plucain, J., T. Hindré, M. Le Gac, O. Tenaillon, S. Cruveiller, C. Médigue, N. Leiby, et al. 2014. Epistasis and allele specificity in the emergence of a stable polymorphism in *Escherichia coli*. Science 343:1366–1369.

Quandt, E. M., J. Gollihar, Z. D. Blount, A. D. Ellington, G. Georgiou, and J. E. Barrick. 2015. Fine-tuning citrate synthase flux potentiates and refines metabolic innovation in the Lenski evolution experiment. eLife 4:e09696.

Rodríguez, J. A., U. M. Marigorta, D. A. Hughes, N. Spataro, E. Bosch, and A. Navarro. 2017. Antagonistic pleiotropy and mutation accumulation influence human senescence and disease. Nature Ecology and Evolution 1:1–5.

Rozen, D. E., D. Schneider, and R. E. Lenski. 2005. Long-term experimental evolution in *Escherichia coli*. XIII. Phylogenetic history of a balanced polymorphism. Journal of Molecular Evolution 61:171–180.

Ruiz, J. A., R. O. Fernández, P. I. Nikel, B. S. Méndez, and M. J. Pettinari. 2006. Dye (Arc) mutants: Insights into an unexplained phenotype and its suppression by the synthesis of poly (3-hydroxybutyrate) in *Escherichia coli* recombinants. FEMS Microbiology Letters 258:55–60.

Salmon, K. A., S. Hung, N. R. Steffen, R. Krupp, P. Baldi, G. W. Hatfield, and R. P. Gunsalus. 2005. Global gene expression profiling in *Escherichia coli* K12: effects of oxygen availability and ArcA. Journal of Biological Chemistry 280:15084–15096.

Saxer, G., M. D. Krepps, E. D. Merkley, C. Ansong, B. L. Deatherage Kaiser, M. T. Valovska, N. Ristic, et al. 2014. Mutations in global regulators lead to metabolic selection during adaptation to complex environments. PLoS Genetics 10:e1004872.

Setlow, P. 2006. Spores of *Bacillus subtilis*: Their resistance to and killing by radiation, heat and chemicals. Journal of Applied Microbiology 101:514–525.

Shabalina, S. A., L. Y. Yampolsky, and A. S. Kondrashov. 1997. Rapid decline of fitness in panmictic populations of *Drosophila melanogaster* maintained under relaxed natural selection. Proceedings of the National Academy of Sciences of the USA 94:13034–13039.

Shannon, P., A. Markiel, O. Ozier, N. S. Baliga, J. T. Wang, D. Ramage, N. Amin, et al. 2003. Cytoscape: A software environment for integrated models of biomolecular interaction networks. Genome Research 13:2498–2504.

Shewaramani, S., T. J. Finn, S. C. Leahy, R. Kassen, P. B. Rainey, and C. D. Moon. 2017. anaerobically grown *Escherichia coli* has an enhanced mutation rate and distinct mutational spectra. PLoS Genetics 13:e1006570.

Siegal, M. L., and J. Leu. 2014. On the nature and evolutionary impact of phenotypic robustness mechanisms. Annual Review of Ecology, Evolution, and Systematics 45:495–517.

Sniegowski, P. D., P. J. Gerrish, and R. E. Lenski. 1997. Evolution of high mutation rates in experimental populations of *E. coli*. Nature 387:703–705.

Sokal, R. R., Rohlf, F. J. 1995. Biometry: The principles and practice of statistics in biological research, 3rd Edition. W. H. Freeman, New York.

Soo, R. M., J. Hemp, D. H. Parks, W. W. Fischer, and P. Hugenholtz. 2017. On the origins of oxygenic photosynthesis and aerobic respiration in Cyanobacteria. Science 355:1436–1440.

Stirling, F., L. Bitzan, S. O’Keefe, E. Redfield, J. W. K. Oliver, J. Way, and P. A. Silver. 2017. Rational design of evolutionarily stable microbial kill switches. Molecular Cell 68:686–697.

Swain, A., and W. F. Fagan. 2019. A mathematical model of the Warburg Effect: Effects of cell size, shape and substrate availability on growth and metabolism in bacteria. Mathematical Biosciences and Engineering 16:168–186.

Szklarczyk, D., J. H. Morris, H. Cook, M. Kuhn, S. Wyder, M. Simonovic, A. Santos, et al. 2017. The STRING database in 2017: Quality-controlled protein-protein association networks, made broadly accessible. Nucleic Acids Research 45:D362–D368.

Tenaillon, O., J. E. Barrick, N. Ribeck, D. E. Deatherage, J. L. Blanchard, A. Dasgupta, G. C. Wu, et al. 2016. Tempo and mode of genome evolution in a 50,000-generation experiment. Nature 536:165–170.

Travisano, M., and R. E. Lenski. 1996. Long-term experimental evolution in *Escherichia coli*. IV. Targets of selection and the specificity of adaptation. Genetics 143:15–26.

Unden, G., and J. Bongaerts. 1997. Alternative respiratory pathways of *Escherichia coli*: energetics and transcriptional regulation in response to electron acceptors. Biochimica et Biophysica Acta 1320:217–234.

Unden, G., S. Becker, J. Bongaerts, J. Schirawski, and S. Six. 1994. Oxygen regulated gene expression in facultatively anaerobic bacteria. Antonie van Leeuwenhoek 66:3–22.

Van den Bergh, B., T. Swings, M. Fauvart, and J. Michiels. 2018. Experimental design, population dynamics, and diversity in microbial experimental evolution. Microbiology and Molecular Biology Reviews 82:e00008–18.

Vasi, F., M. Travisano, and R. E. Lenski. 1994. Long-term experimental evolution in *Escherichia coli*. II. Changes in life-history traits during adaptation to a seasonal environment. American Naturalist 144:432–456.

Wagner, G. P., and J. Mezey. 2000. Modeling the evolution of genetic architecture: A continuum of alleles model with pairwise AxA epistasis. Journal of Theoretical Biology 203:163–175.

Warburg, O. 1956. On the origin of cancer cells. Science 123:309–314.

Wielgoss, S., J. E. Barrick, O. Tenaillon, M. J. Wiser, W. J. Dittmar, S. Cruveiller, B. Chane-Woon-Ming, et al. 2013. Mutation rate dynamics in a bacterial population reflect tension between adaptation and genetic load. Proceedings of the National Academy of Sciences of the USA 110:222–227.

Williams, G. C. 1957. Pleiotropy, natural selection, and the evolution of senescence. Evolution 11:398–411.

Wiser, M. J., N. Ribeck, and R. E. Lenski. 2013. Long-term dynamics of adaptation in asexual populations. Science 342:1364–1367.

Woods, R., D. Schneider, C. L. Winkworth, M. A. Riley, and R. E. Lenski. 2006. Tests of parallel molecular evolution in a long-term experiment with *Escherichia coli*. Proceedings of the National Academy of Sciences of the USA 103:9107–9112.

